# Dendrites of DG granule cells contribute to pattern separation by controlling sparsity

**DOI:** 10.1101/067389

**Authors:** Spyridon Chavlis, Panagiotis C. Petrantonakis, Panayiota Poirazi

## Abstract

The hippocampus plays a key role in pattern separation, the process of transforming similar incoming information to highly dissimilar, non-overlapping representations. Sparse firing granule cells (GCs) in the dentate gyrus (DG) have been proposed to undertake this computation, but little is known about which of their properties influence pattern separation. Dendritic atrophy has been reported in diseases associated with pattern separation deficits, suggesting a possible role for dendrites in this phenomenon. To investigate whether and how the dendrites of GCs contribute to pattern separation, we build a simplified, biologically relevant, computational model of the DG. Our model suggests that the presence of GC dendrites is associated with high pattern separation efficiency while their atrophy leads to increased excitability and performance impairments. These impairments can be rescued by restoring GC sparsity to control levels through various manipulations. We predict that dendrites contribute to pattern separation as a mechanism for controlling sparsity.

## Introduction

The hippocampus is known to be involved in memory formation, storage and consolidation (Squire et al., 2004), but its specific functionalities remain a mystery. One such functionality is the ability to rapidly store non-overlapping representations of similar inputs and thereafter, retrieve them given a partial or noisy stimulus. Theoretical models refer to those processes as pattern separation and pattern completion, respectively (Marr, 1971; Treves and Rolls, 1994; Yassa and Stark, 2011; Santoro, 2013). The Dentate Gyrus (DG), in particular, has been proposed to implement pattern separation by sparsifying and orthogonalizing its input, coming mainly from the Entorhinal Cortex (EC), and thereafter, projecting this information to the CA3 area via the mossy fibers (Treves and Rolls, 1994). DG has been hypothesized to separate two distinct but overlapping patterns through the activation of different Granule Cells (GCs), through the expression of different firing rates in identical neuronal populations (Deng et al., 2010) or a combination of the two. While several studies have investigated pattern separation both in rodents (Leutgeb et al., 2004, 2005, 2007) and humans (Kirwan and Stark, 2007; Bakker et al., 2008; Lacy et al., 2011; Motley and Kirwan, 2012), the role of dendrites in this phenomenon remains unknown.

The DG is the first subregion of the hippocampus that receives incoming information from other brain areas. DG principal neurons, the Granule Cells (GCs), receive input from excitatory afferents coming from EC layer II cells and project to the CA3 subregion. In addition, they receive input from other DG excitatory cells, the Mossy Cells (MCs), and various interneurons (Sik et al., 1997) with Basket Cells (BCs) and Hilar Perforant Path associated (HIPP) cells being the most important. MCs form an inhibitory circuit as their axons contact the BCs. The net effect of MC excitation to both GCs and BCs is considered to be inhibitory (Jinde et al., 2012).

Experimental studies have shown that only a small population of GCs, ~5%, are active in a single context (Marrone et al., 2011; Satvat et al., 2011; Danielson et al., 2016), a phenomenon termed sparse coding (Olshausen and Field, 2004). It has been proposed that sparse coding in GCs enhances pattern separation by recruiting different subgroups of GCs to encode similar incoming stimuli (Treves et al., 2008; Petrantonakis and Poirazi, 2014, 2015). Computational models (Santhakumar et al., 2005; Yim et al., 2015) and experimental studies (Nitz and McNaughton, 2004; Jinde et al., 2012) have proposed that inhibition controls GC activity which, in turn, mediates pattern separation (Myers and Scharfman, 2009, 2011; Ikrar et al., 2013; Faghihi and Moustafa, 2015).

The ability to perform pattern separation is critical for normal brain functioning and its impairment is associated with cognitive decline. Diseases such as schizophrenia (Das et al., 2014) and Alzheimer’s Disease (AD) (Ally et al., 2013), where cognitive decline is evident, are both characterized by pattern separation deficiencies. Interestingly, these conditions are also characterized by alterations in the anatomical properties of GC dendrites, such as a decrease in the total dendritic length (Einstein et al., 1994) and spine loss (Jain et al., 2012). Dendritic growth on the other hand has been associated with pattern separation enhancements. Specifically, voluntary running was recently shown to enhance pattern separation and this enhancement was attributed to an increase in the neurogenesis rate that was accompanied by increased GC dendrite outgrowth in active compared to sedentary animals (Bolz et al., 2015). These findings suggest that the dendrites of GCs are likely to play a key role in pattern separation mediated by the DG.

To investigate this possibility, we implemented a morphologically simple, yet biologically relevant, scaled-down spiking neural network model of the DG. The model consists of four types of cells (MCs, BCs, HIPP and GCs) modeled as simplified integrate-and-fire neurons. The GC model alone was extended to incorporate dendrites. The electrophysiological properties of all cell types were calibrated according to a range of experimental data. An advantage of using such a simplified approach lies in the small number of parameters that make it possible to characterize their role in the behavior of the model. Despite its simplicity, the model exhibits realistic pattern separation under several conditions and explains how inhibition to GCs provided directly from BCs and indirectly via the inhibitory circuitry through MCs impact this task, as suggested by a number of recent studies (Myers and Scharfman, 2011; Jinde et al., 2012), thus, supporting its biological relevance. We use the model to investigate whether and how GC dendrites may contribute to pattern separation.

## Results

### The Dentate Gyrus model

The DG network model consists of four different neuronal types, namely GCs, BCs, MCs, and HIPP cells. MCs, BCs and HIPP cells are modeled as adaptive exponential I&F (aEIF) point neurons (Brette and Gerstner, 2005). GCs consist of a leaky integrate-and-fire somatic compartment connected to a variable number of active dendritic compartments (3, 6 or 12) whose morphology relies loosely on anatomical data (see Materials and Methods). I&F models were selected primarily because of their correspondence with experimental parameters (e.g. the C_m_, R_m_, R_in_) that facilitated constraining with experimental data. Since dentate GCs are known to have 10–15 dendrites (Claiborne et al., 1990), we consider the GC model with 12 dendrites as the control. A schematic illustration of the DG network is shown in Figure 1A.

**Figure 1.**
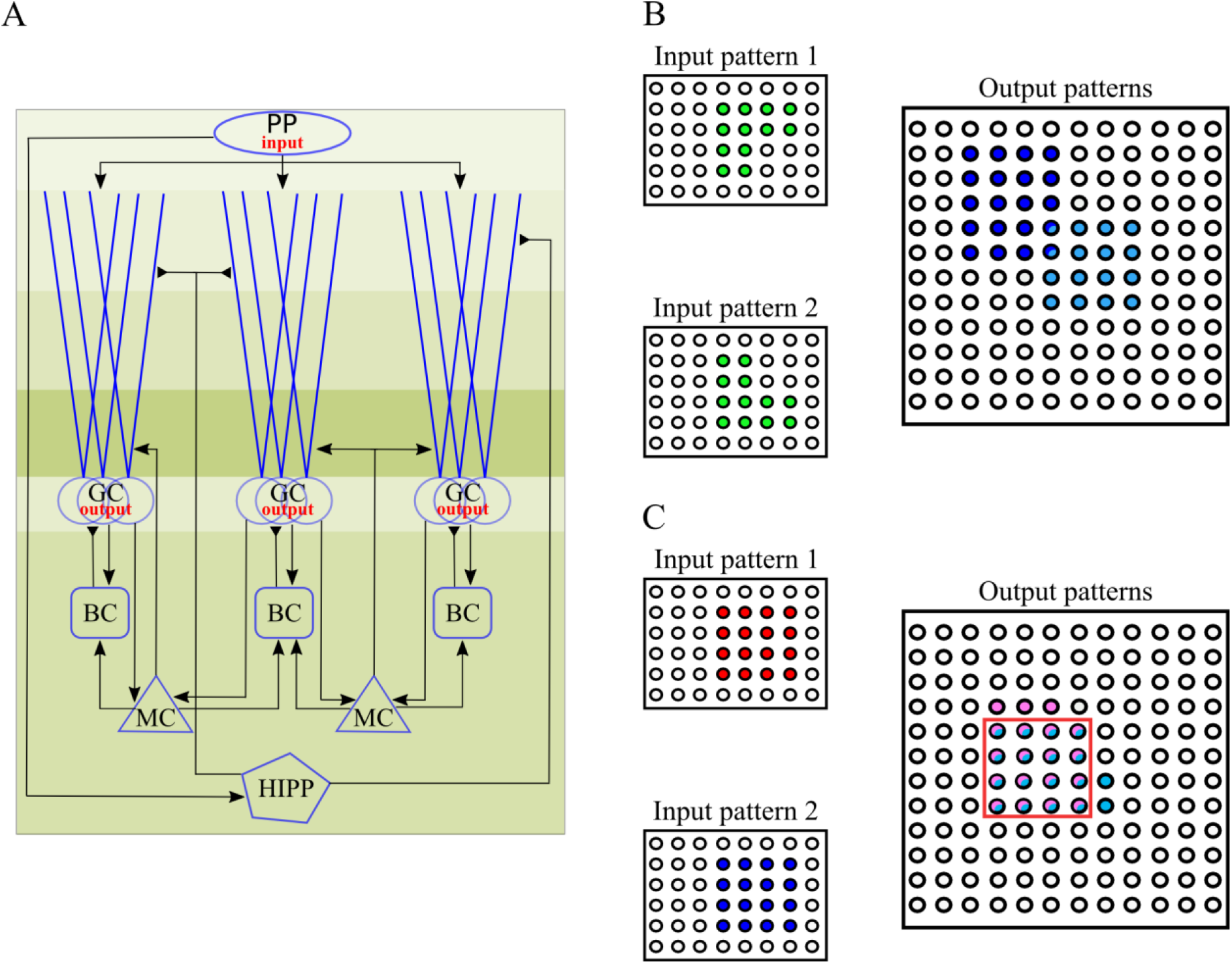
**A**. Schematic representation of the implemented DG network model. Different shades of green illustrate the layer division. PP: Perforant Path, GC: Granule Cells, BC: Basket Cells, MC: Mossy Cells, HIPP: Hilar Perforant Path-associated cells. Perforant Path afferents curry the input to the network, and project on both the GCs and the HIPP cells. MCs and GCs are connected in a recurrent manner. MCs also excite the BCs. Inhibition of GCs is provided through BCs and HIPP cells directly and indirectly via MCs. Note that HIPP cells contact the distal dendrites rather than the soma of GCs. GCs provide the output of the DG network. **B**. Schematic representation of pattern separation using population-based coding. When two highly overlapping EC inputs (input 1 & 2, with identical mean firing rates) arrive in DG, the corresponding outputs are highly dissimilar. Note that the output pattern is sparse because of the low number of GCs that encode any given pattern. **C**. Schematic representation of pattern separation using rate-based coding. When two highly overlapping EC inputs (input 1 & 2, with different mean firing rates but identical input populations) arrive in DG, the corresponding outputs are highly dissimilar in their firing rates but also likely to differ in the populations they activate. Rate distances are estimated over the set of common neurons (red box).

All computational neuron models were validated against experimental data with respect to their activity and basic electrophysiological properties (Lübke et al., 1998; Bartos et al., 2001; Krueppel et al., 2011). The spiking profiles of the four neuronal model types are depicted in Figure 2 while the respective I-V and I-F curves compared against experimental data are shown in Figure 2-figure supplement 1 and Figure 2-figure supplement 2. The control (12-dendrite) GC model was validated against the experimental data of (Krueppel et al., 2011) with respect to its dendritic input-output function, which was found to be slightly above linearity (see Figure 2-figure supplement 3(A)). Table 1 lists the model and experimental values of basic intrinsic properties for each of these cell types. Overall, the electrophysiological properties of the computational models of neurons included in the DG network are in fair alignment with experimental findings. It should be noted that spiking profiles are not taken into account when estimating pattern separation in the DG network model. A GC model neuron was considered active even if it produced a single spike. Thus, we chose to fit average values rather than temporal profiles of the model neurons.

**Figure 2.**
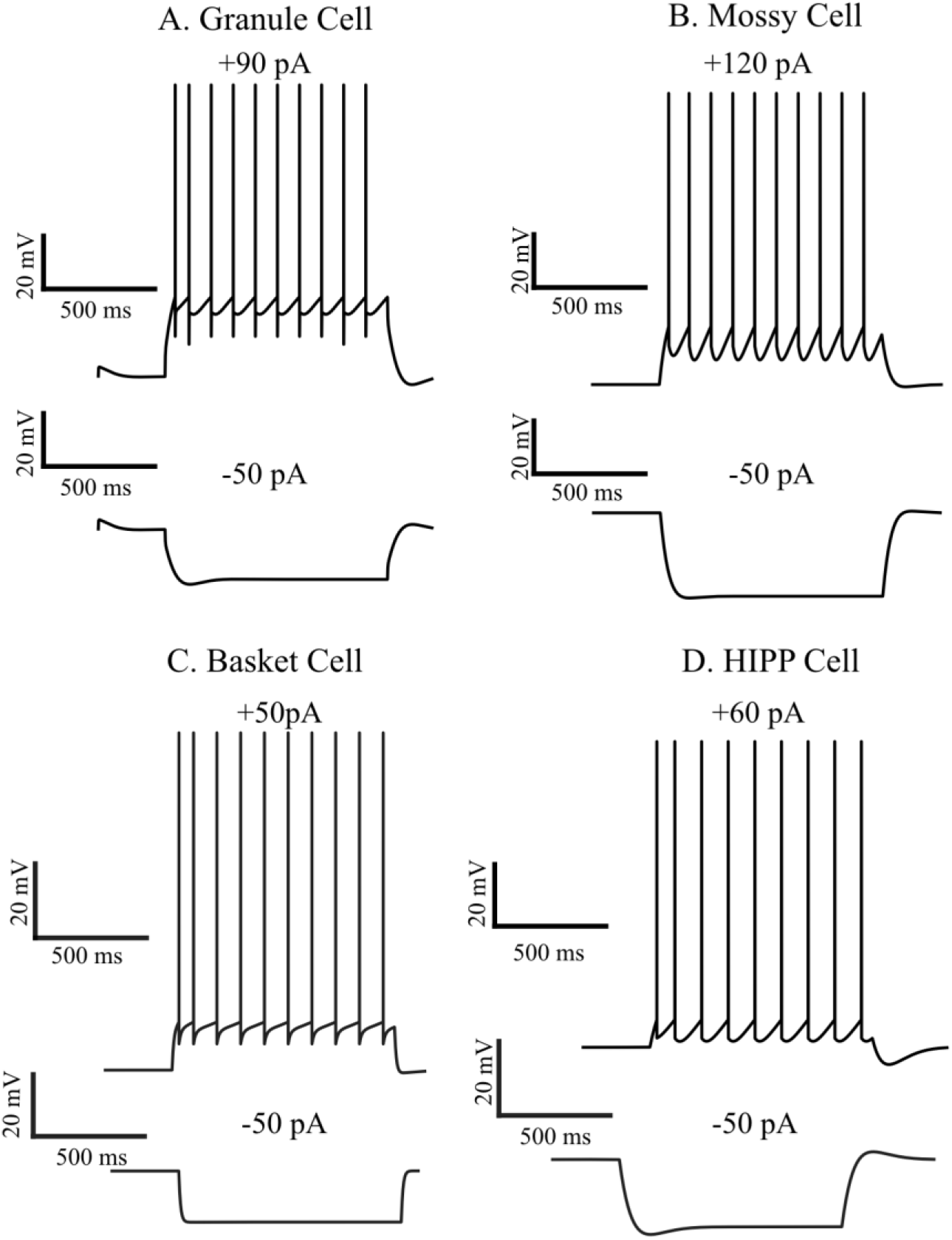
Firing traces of the four model cells in response to current injection (1 second). Note that APs are not explicitly modeled in I&F neurons. **A**: The somatic membrane voltage of the granule cell (GC) model in response to +90 pA (top), and −50 pA (bottom) somatic current injections. **B**. Same as in **A**, for the mossy cell (MC) with +300 pA (top) and −300pA (bottom) current injections. **C**. Same as in **A**, for the Basket cell (BC), with +200 pA (top) and −50 pA (bottom) current injections. **D**. Same as in **A**, for the HIPP cell, with +300 pA (top) and −50 pA (bottom) current injections. Current – Voltage (I-V) curves are given in Figure 2-figure supplement 1, while Current – frequency plots are shown in Figure 2-figure supplement 2. Moreover, Figure 2-figure supplement 3 shows the somatic EPSP against Arithmetic sum for GCs under control, pruning and growth conditions. In Figure 2-figure supplement 4 and 5, the corresponding EPSP vs. Arithmetic sum curves for GCs after matching Rin and sparsity are given, respectively.

**Table 1.** Physiological properties of real and model neurons

| Cell type | Input resistance $R_{in}$ (MΩ) | | Membrane time constant, $\tau_m$ (ms) | | Sag ratio | | Maximum firing rate (Hz) | |
| --- | --- | --- | --- | --- | --- | --- | --- | --- |
|  | Model | Biological | Model | Biological | Model | Biological | Model | Biological |
| Granule cells | 360 | 292 ± 34 | 41.2 | 31 ± 2 | 0.91 | 0.96 ± .1 | 60 | 70 ± 10 |
| Mossy cells | 105 | 199 ± 19 | 33.7 | 35 ± 5 | 0.98 | 0.81 ± .3 | 45 | 50 ± 6 |
| Basket cells | 55 | 56 ± 9 | 9.67 | 10 ± 1 | 0.99 | 0.97 ± .02 | 247 | 230 ± 15 |
| HIPP cells | 363 | 371 ± 47 | 21.4 | 15 ± 0 | 0.84 | 0.82 ± .04 | 113 | 101 ± 24 |
| Biological value | Lubke et al., 1998,<br>Krueppel et al., 2009 |  | Lubke et al., 1997, Ratzliff<br>et al., 2004 |  | Lubke et al., 1998, Bartos<br>et al., 2001 |  | Lubke et al., 1997 |  |

### Pattern separation in the DG model

Following validation of the individual neuron models, we tested the DG network’s ability to implement pattern separation when presented with pairs of inputs characterized by various degrees of similarity (modeled as overlap in the two activated EC populations). An example of two such input patterns and their corresponding output is schematically illustrated in Figure 1B. The network model is deemed capable of separating similar input patterns if the value of a distance metric *f* is substantially larger in the DG output (GC activity) compared to its input (EC cell activity). Pattern separation is primarily estimated by looking at the differences in the populations of neurons that encode each input (‘population distance’, *f*_1_). To account for the possibility of rate-based pattern separation (Deng et al., 2010), we also simulate the network under conditions where the two input pairs are identical in terms of EC ‘population distance’ (i.e., *f*_1(*input*)_ = 0) but differ in their mean firing rates (graphically depicted in Figure 1C). Pattern separation in this case is measured by looking at differences in the average firing rate of the neurons encoding both inputs (‘rate distance’, *f*_2_). The ‘rate distance’ is estimated both at the input (EC cells) and the output (GCs) levels (see Materials and Methods). Pattern separation is successfully performed if the ‘rate distance’ of the output (*f*_2_(*output*)) is larger than that of the input (*f*_2(*input*)_). The network’s ability to perform pattern separation across all (200) input pairs measured with both metrics is shown in Table 2. These results demonstrate that two inputs that are quite similar in their topology (i.e., originate largely (*f*_1_) or entirely from the same neurons, perhaps with different firing rates (*f*_2_)), induce substantially different activation patterns of the GCs that they impinge on (outputs) in the DG network.

**Table 2.** DG network (12-dendrite model) performance on a simple pattern separation task.

| Input Similarity ($f_{1(input)}$) | Output Similarity ($f_{1(output)}$) mean ± sem |
| --- | --- |
| 0.10 | 0.43 ± 0.007 |
| 0.20 | 0.52 ± 0.007 |
| 0.30 | 0.59 ± 0.008 |
| 0.40 | 0.64 ± 0.008 |
| Input Similarity ($f_{2(input)}$) mean ± sem | Output Similarity ($f_{2(output)}$) mean ± sem |
| 0.16 ± 0.05 | 0.66 ± 0.01 |

### Understanding the role of Inhibition in Pattern separation

After establishing the network’s ability to perform pattern separation, we tested its validity against experimental data with respect to the role of inhibition in this phenomenon. The network model reproduced the recent findings of Engin et al., whereby inhibition exerted by BCs was critical for the sparse firing of GCs (Engin et al., 2015). Specifically, removal of all BCs resulted in an over-excitation of the GC model population (more than 50% of GCs responded strongly to any input). This over-excitation in turn impaired pattern separation, as the GC populations responding to the two inputs became nearly identical and fired at very similar, high frequencies. These findings are also in line with experimental evidence reporting increased memory interference under conditions of reduced BC activity (Engin et al., 2015).

Since MCs have also been suggested to control the excitability of DG granule cells (Jinde et al., 2013), we simulated a complete MC-loss lesion (Figure 3A) as per (Ratzliff et al., 2004). This manipulation led to an increase in the proportion of active GCs for all input patterns tested (Figure 3B, 3D), and a decrease in pattern separation efficiency, measured either with the population (*f*_1_, Figure 3C) (Wilcoxon test, *W_1_=2095.5*, *p_1_≈10^−9^, 95% Cl_1_ [0.05, 0.09], W_2_=2242.0, p_2_≈10^−11^, 95% Cl_2_ [0.06, 0.10], W_3_=2363.0, p_3_≈10^−11^, 95% Cl_3_ [0.07, 0.11], W_4_=2444.0, p_4_≈10^−16^, 95% Cl_4_ [0.08, 0.12]*) or rate metrics (*f*_2_, Figure 3E) (Wilcoxon test, *W=930.0, p=0.0276, 95% Cl [0.01, 0.07]*). The subscripts (form one to four) in the Wilcoxon test statistic (W), p-value, confidence interval (CI) correspond to the four experiments with different input overlaps (see Materials and Methods). The observed decrease in sparsity under the MC-loss condition was accompanied by small increases in the excitability levels of GC models. Specifically, for the population coding experiment, the mean GC firing frequency increased from 3.5 Hz to 4.82 Hz, while for the rate-based coding experiment from 3.75 Hz to 4.94 Hz for 40 Hz inputs and from 5.24 Hz to 8.07 Hz for 50 Hz inputs. These findings are in line with the experimental data of (Ratzliff et al., 2004) where MC-loss did not lead to an over-excitation of GCs.

**Figure 3.**
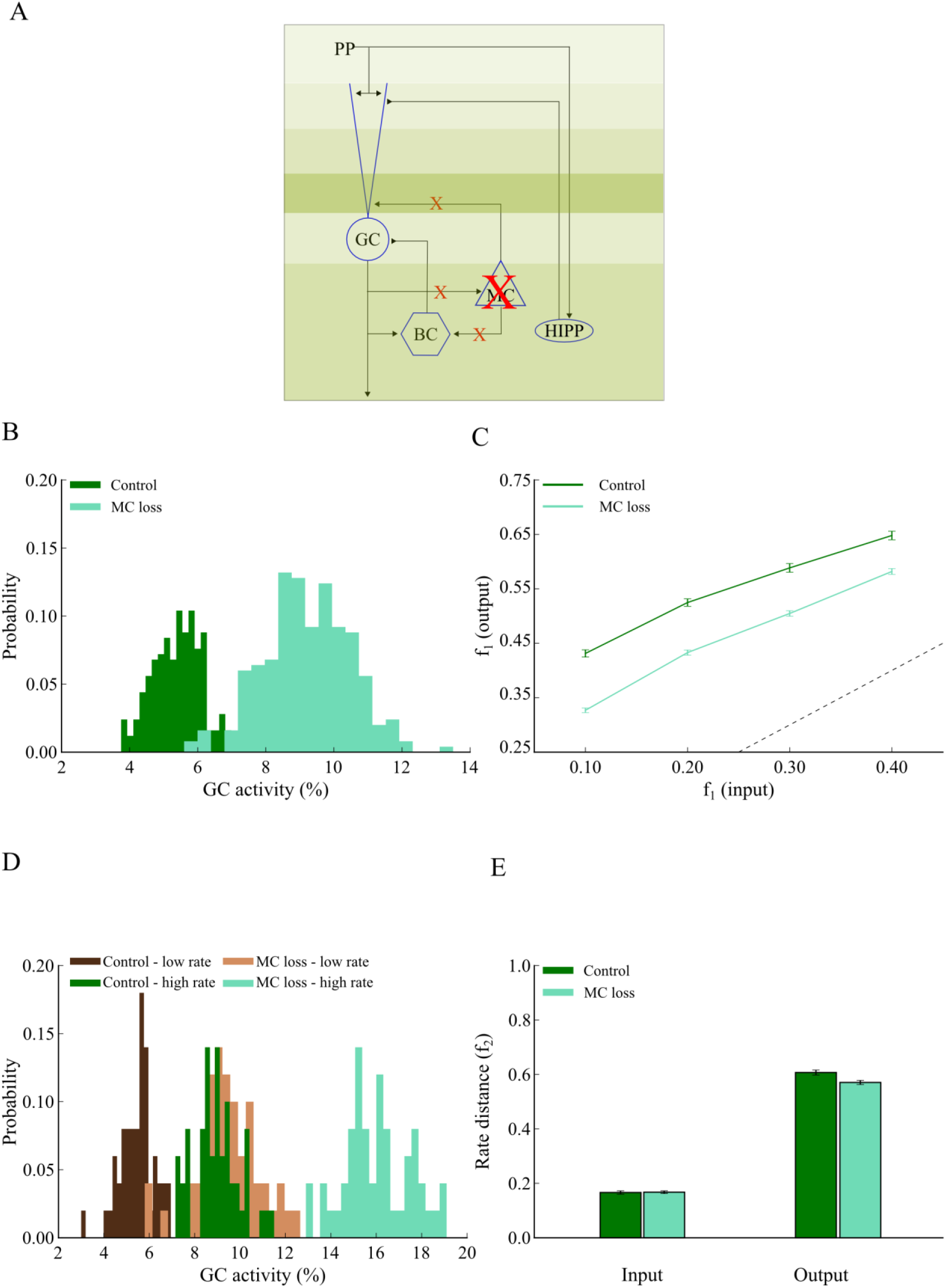
Complete mossy cell loss reduces pattern separation efficiency in the DG network. **A**. Schematic representation of the 12-dendrite DG network with MC loss. **B**. The corresponding probability density functions of GC activity in response to 40 Hz input for the control (dark green) and MC loss conditions (light green). The mean activity is 5% and 9% for the control and MC loss networks, respectively. The respective mean firing rates of GCs are shown in Table 3. The histograms were calculated with 20 bins each. **C**. Input / Output population distances (*f*_1_) for the control (dark green) and MC loss (light green) networks estimated using input patterns with increasing similarity. The dashed line denotes the limit above which the model performs pattern separation. MC loss reduces pattern separation efficiency for all input patterns tested. Error bars represent the standard error of the mean (Erevitt and Skrondal, 2010). **D**. Probability density functions of GC activity using control and MC loss models presented with two input patterns that differ only in their firing rates. Shades of green represent the high frequency input (50 Hz), while shades of brown represent the low frequency input (40Hz). Dark and light shades represent the control and MC loss condition, respectively. The respective mean firing rates of GCs are shown in Table 3. **E**. Input / Output rate distances (*f*_2_) for the control (dark green) and MC loss (light green) networks estimated using two input patterns with different firing frequencies, 40 and 50 Hz respectively. MC loss slightly reduces the efficiency of pattern separation.

Taken together, the proposed DG network model a) exhibits single-neuron response properties that are in good agreement with experimental data, b) implements a connectivity profile that relies on experimental observations, c) exhibits robust pattern separation and d) replicates experimental data about the role of BCs and MC cells in the aforementioned task. These features support the biological plausibility of the model which is next used to investigate how dendrites contribute to pattern separation.

### Dendrites and Pattern separation

#### Dendritic pruning

The main question of interest in this work is whether and how dendrites may contribute to pattern separation. To answer this question we started by examining whether the number of dendrites correlates with pattern separation performance as we prune the GC (sister) branches from 12 (control) to 6 and 3 (Figure 4A). To assess the effect of dendritic number, we kept all other parameters (maximum dendritic length, dendritic diameter in the IML, MML and OML layers, membrane capacitance, “leak” conductance, number of activated synapses and input firing rates) of the three GC models identical to those of the control. Dendritic integration properties for the pruned GC models are shown in Figure 2-figure supplement 3 (B). The control DG network (12-dendrite model) was calibrated to have a mean population sparsity level of 2–5% as per (Treves et al., 2008; Danielson et al., 2016), meaning that only a small fraction of GCs were active for any given stimulus.

**Figure 4.**
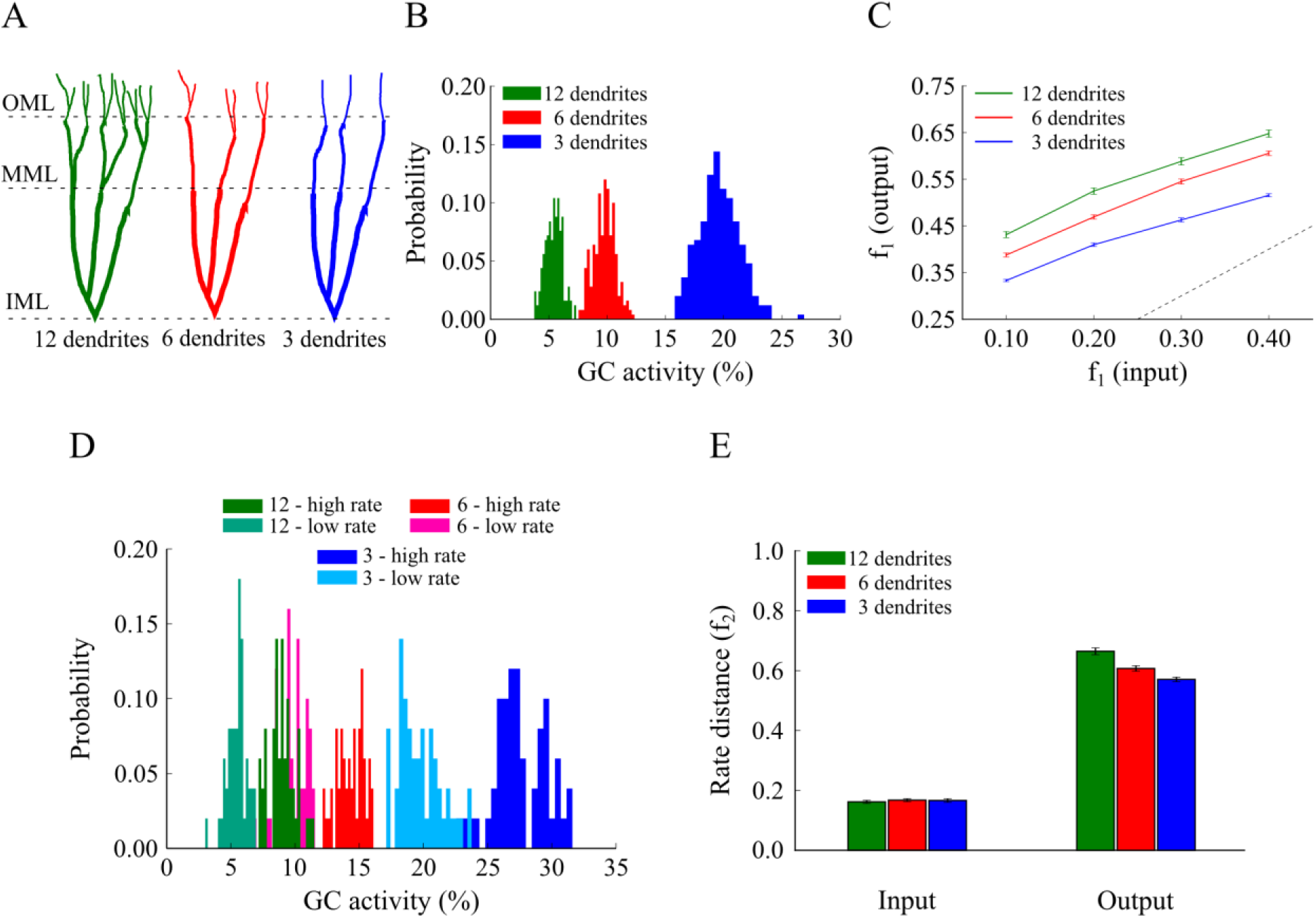
Effect of GC dendrite pruning on pattern separation. **A**. A schematic drawing of the three GC models with 12, 6 and 3 dendrites. **B**. Corresponding probability density functions of GC activity for the three GC models in response to a single input pattern at 40 Hz. The mean activity is inversely analogous to the number of dendrites: as the number of dendrites decreases, the GC population becomes more active, i.e., less sparse. The respective mean firing rates of GCs are shown in Table 3. **C**. Input / Output population distances (*f*_1_), for the 12-dendrite (green), 6-dendrite (red) and 3-dendrite (blue) GC models in response to the presentation of two overlapping input patterns at 40 Hz. Input pairs were generated so as to have decreasing amounts of overlap, as depicted in Figure 1B. The dashed line denotes the limit above which the model performs pattern separation. Performance declines as the dendritic tree has fewer dendrites. Error bars represent the standard error of the mean across trials. **D**. Probability density functions of GC activity for the three GC models in response to presentation of two input patterns with different firing rates (low rate = 40 Hz, high rate = 50 Hz), as depicted in Figure 1C. GC activity increases with the number of dendrites and the input firing rate. The respective mean firing rates of GCs are shown in Table 3. **E**. Input / Output rate distances (i.e., *f*_2_) for the three models. All three models perform pattern separation, as the rate distances in the input are significantly higher than the corresponding ones in the output.

Interestingly, pattern separation of pairs of inputs with increasing similarity (measured either by population distances, Figure 4C, or by rate distances, Figure 4E) was successfully performed in the control as well as both pruned models for all pairs of inputs tested. The efficiency of pattern separation however correlated with the number of dendrites in GCs (Figure 4C, 4E), with the 12-dendrite GC model achieving the best performance for both population (12-dendrite vs. 6-dendrite: Wilcoxon test, *W_1_=1849.0, p_1_≈10^−5^, 95% Cl_1_ [0.02, 0.06], W_2_=1818.5, p_2_≈10^−4^, 95% Cl_2_ [0.02, 0.06], W_3_=2060.5, p_3_≈10^−8^, 95% Cl_3_ [0.04, 0.07], W_4_=1948.0, p_4_≈10^−6^ 95% Cl_4_ [0.03, 0.06]* and 12-dendrite vs. 3-dendrite: Wilcoxon test, *W_1_=2471.0, p_1_≈10^−16^, 95% Cl_1_ [0.11, 0.15], W_2_=2462.0, p_2_≈10^−16^, 95% Cl_2_ [0.11, 0.14], W_3_=2473.0, p_3_≈10^−16^, 95% Cl_3_ [0.10, 0.13], W4=2451.0, *p*_4_≈10^−16^, 95% Cl_4_ [0.08, 0.11]*) and rate based-metrics (12-dendrite vs. 6-dendrite: Wilcoxon test, *W=1764.0, p=0.0004, 95% Cl [0.03, 0.09]* and 12-dendrite vs. 3-dendrite: Wilcoxon test, *W=2072.0, p≈10^−8^, 95% Cl [0.06, 0.12]*) (i.e., the highest *f*_1(*outPut*)_ and *f*_2(*output*)_, respectively).

These findings are better understood by looking at the sparsity levels exhibited by the three GC network models. As shown in Figure 4B, the percentage of active GCs for the population-based experiment increased substantially when the number of dendrites was reduced. It rose from ~5%, to ~10% and ~20%, for GC model cells with 12, 6 and 3 dendrites, respectively. These differences in activity distributions were statistically significant (12-dendrite vs. 6-dendrite: Kolmogorov-Smirnov test, *D=1.000, p≈10^−16^* and 12-dendrite vs. 3-dendrite: Kolmogorov-Smirnov test, *D=1.000. p≈10^−16^*). The activity levels of the three models followed a similar pattern in the case of rate-based coding (Figure 4D); Input patterns with high firing rates induced lower sparsity levels in the 12-dendrite GC model, followed by the 6-dendrite model and then the 3-dendrite model (12-dendrite vs. 6-dendrite: Kolmogorov-Smirnov test, *D=1.000, p≈10^−16^* and 12-dendrite vs. 3-dendrite: Kolmogorov-Smirnov test, *D=1.000. p≈10^−16^*). These data reveal that high levels of sparsity (namely low levels of GC activity) and pattern separation efficiency in the network model are a direct consequence of having multiple dendrites and suggest that dendrites may contribute to pattern separation through their effects on sparsity.

### Dendritic growth

To further test whether the presence of dendrites helps pattern separation by increasing sparsity, we also simulated the opposite process, namely the growth of dendrites. We built GC models with 3, 6 and 12 dendrites with shapes that roughly mimic the stages of dendritic growth (Figure 5A): starting with a GC model consisting of 3 thick dendritic compartments and adding a branch point with two thinner sister branches at each terminal dendrite we end up with a 12-dendrite model which is identical with our control one (see Materials and Methods). Dendritic integration properties for the growth GC models are shown in Figure 2-figure supplement 3 (C). Note that in this simulation, the number, length and mean diameter of dendrites differ between the three models.

**Figure 5.**
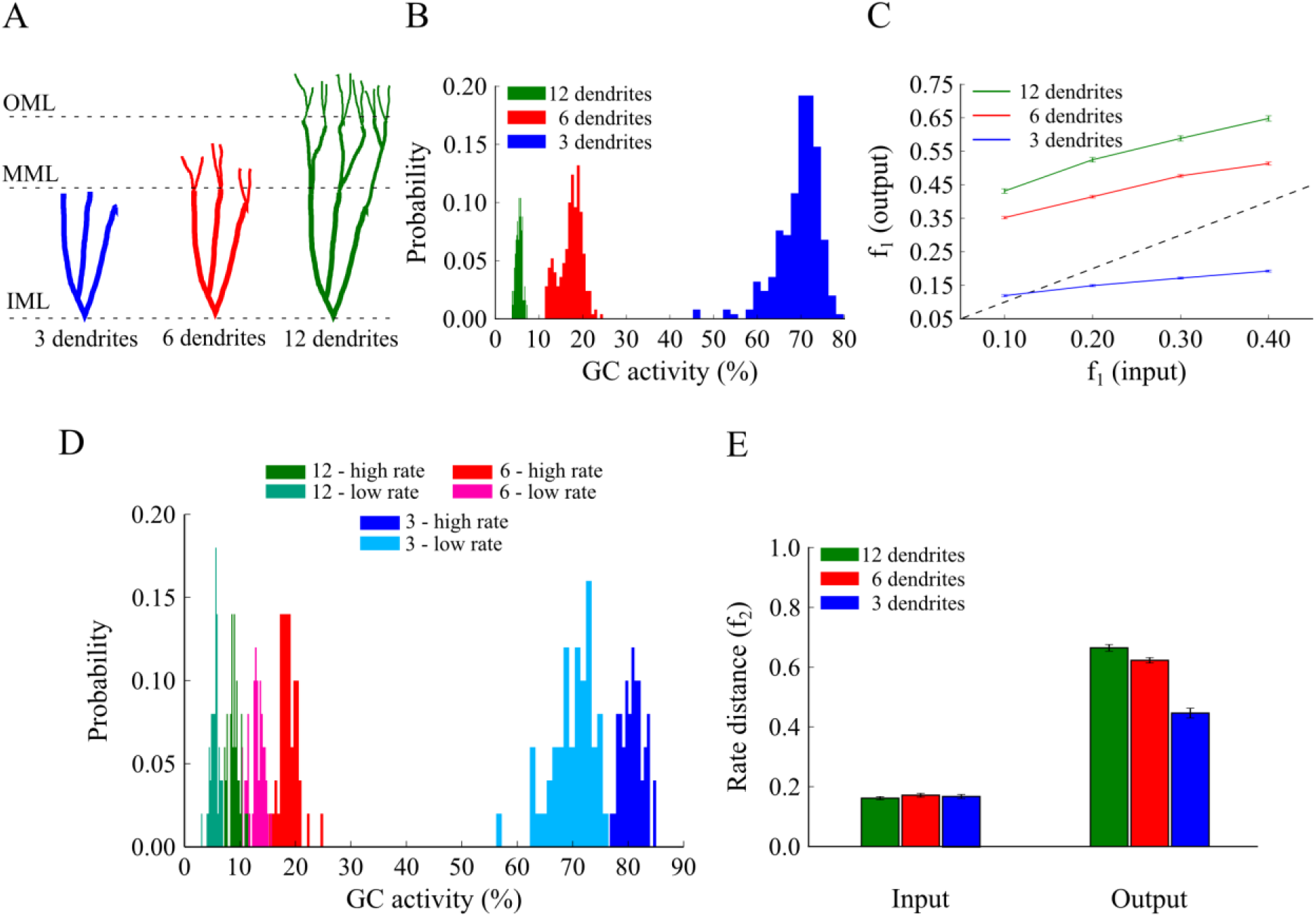
Effect of GC dendrite growth on pattern separation. **A**. A schematic drawing of the three GC models with 3, 6 and 12 dendrites. **B**. Corresponding probability density functions of GC activity for the three GC models in response to a single input pattern at 40 Hz. The mean activity is inversely analogous to the number/length of dendrites: as dendrites grow, the GC population becomes sparser, i.e., fewer GCs are active. The respective mean firing rates of GCs are shown in Table 3. **C**. Input / Output population distances (*f*_1_), for the 3-dendrite (blue), 6-dendrite (red) and 12-dendrite (green) GC models in response to the presentation of two overlapping input patterns at 40 Hz, with different degrees of overlap, as depicted in Figure 1B. The dashed line denotes the limit above which the model performs pattern separation. Performance improves with the growth of dendrites. **D**. Probability density functions of GC activity for the three GC models in response to presentation of two input patters with different firing rates (low rate = 40 Hz, high rate = 50 Hz), as depicted in Figure 1C. GC activity decreases with the number of dendrites and increases with the input firing rate. The respective mean firing rates of GCs are shown in Table 3. E. Input / Output rate distances (i.e., *f*_2_) for the three models. The number of GC dendrites is again analogous to the pattern separation performance.

In line with the findings of the pruning experiment, the percentage of active GCs declined as the number of dendrites increased (Figure 5B,D). The average GC activity for the population-coding experiment started at ~70% for the network with 3 GC dendrites, dropped to ~28% for the network with 6 GC dendrites and to ~5% for the 12-dendrite (control) model (all differences were statistically significant, (12-dendrite vs. 6-dendrite: Kolmogorov-Smirnov test, *D=1.000, p≈10^−16^* and 12-dendrite vs. 3-dendrite: Kolmogorov-Smirnov test, *D=1.000. p≈10^−16^*). Similar differences in GC sparsity were seen in the rate-based experiment (Figure 5D). In both cases, the 3-dendrite model exhibited much higher activity levels than the ones seen in the pruning experiment (Figure 4B,D), primarily because of additional alterations in dendritic length and diameter. Moreover, pattern separation measured by the population metric was completely impaired in the 3-dendrite model (Figure 5C, blue line falls below the diagonal) while performance based on rate coding remained above baseline (Figure 5E, blue bar). Pattern separation was successfully performed in the 6-dendrite model albeit with lower efficiency compared to the 12-dendrite model, as evaluated both with the population (12-dendrite vs. 6-dendrite: Wilcoxon test, *W_1_=2456.0, p_1_≈10^−16^, 95% Cl_1_ [0.12, 0.15], W_2_=2421.0, p_2_≈10^−15^, 95% Cl_2_ [0.09, 0.13], W_3_=2466.0, p_3_≈10^−16^, 95% Cl_3_ [0.09, 0.13], W_4_=2368.0, p_4_≈10^−14^, 95% Cl_4_ [0.06, 0.09]* and 12-dendrite vs. 3-dendrite: Wilcoxon test, *W_1_. 4=2500.0, p_1–4_≈10^−16^, 95% Cl_1_ [0.44, 0.48], 95% Cl_2_ [0.40, 0.43], 95% Cl_3_ [0.36, 0.39], 95% Cl_4_ [0.29, 0.32]*,) and rate distance metrics (12-dendrite vs. 6-dendrite: Wilcoxon test, *W=1638.0, p=0.0076, 95% Cl [0.01, 0.07]* and 12-dendrite vs. 3-dendrite: Wilcoxon test, *W=2305.0, p≈10^−13^, 95% Cl[0.17, 0.25])* (Figure 5C,E).

Taken together, the “pruning” and “growth” simulations suggest a strong link between pattern separation efficiency and GC population sparsity (Aimone et al., 2011, Deng et al., 2010) and predict that dendrites may serve as a mechanism for increasing the sparsity of the GC population which in turn enhances pattern separation.

### Controlling sparsity with non-dendritic mechanisms

The above simulations predict that dendrites can be sufficient for mediating sparsity, which in turn enhances pattern separation. We next ask whether they are necessary for this task, as it is possible that a GC neuron can counteract the decrease in sparsity induced by having fewer/shorter dendrites through alternative mechanisms.

#### *Input resistance* does *not explain differences in sparsity or pattern separation efficiency*

First, we assessed the effect of input resistance differences among the various GC models on sparsity and pattern separation efficiency. A possible explanation of the above findings is that a GC model with a small dendritic tree has increased input resistance (as can be seen in Supplementary file 1A) which in turn leads to higher excitability and decreased sparsity. We thus corrected the input resistance in the 6- and 3-dendrite models to match the one in the 12-dendrite model (at the soma) by modifying a) the “leak” channel conductance (g_leak_) or b) the size of the somatic compartments (see Supplementary file 1B for the corrected values and Figure 2-figure supplement 4 for the corrected dendritic integration profiles). Specifically, for the pruning models, g_leak_ increased by *1.230* and *1.635* for the 6- and 3-dendrite models while for the growth models, these numbers were *1.596* and *2.438*, respectively. Figure 6 shows the outcome of this correction for both pruning (A-B) and growth (C-D) cases. As evident from the figure, correcting the input resistance by increasing the “leak” conductance in the pruning case, reduced but did not eliminate differences in sparsity (Figure 6A) (12-dendrite vs. 6-dendrite: Kolmogorov-Smirnov test, *D=0.868, p≈10^−16^* and 12-dendrite vs. 3-dendrite: Kolmogorov-Smirnov test, *D=1.000. p≈10^−16^*) or pattern separation efficiency (Figure 6B) (12-dendrite vs. 6-dendrite: Wilcoxon test, *W_1_=1636.5, p_1_=0.0078, 95% Cl_1_ [0.01, 0.07], W_2_=1606.5, p_2_=0.0141, 95% Cl_1_ [0.01, 0.04], W_3_=1732.5, p_3_=0.0009, 95% Cl_1_ [0.02, 0.05], W_4_=1668.5, p_4_=0.0040 95% Cl_4_ [0.01, 0.04]* and 12-dendrite vs. 3-dendrite: Wilcoxon test, *W_1_=2164.0, p_1_≈10^−10^, 95% Cl_1_ [0.06, 0.09], W_2_=2177.5, p_2_≈10^−10^, 95% C1_2_ [0.05, 0.09], W_3_=2284.0, p_3_≈10^−12^, 95% Cl_3_ [0.06, 0.09], W_4_=2151.0, p_4_≈10^−10^ 95% Cl_4_ [0.04, 0.07]*). Similar findings were seen in the growth case, both for the sparsity (Figure 6C) (12-dendrite vs. 6-dendrite: Kolmogorov-Smirnov test, *D=0.956, p≈10^−16^* and 12-dendrite vs. 3-dendrite: Kolmogorov-Smirnov test, *D=1.000, p≈10^−16^*) and pattern separation efficiency (Figure 6D) (12-dendrite vs. 6-dendrite: Wilcoxon test, *W_1_=2252.5, p_1_≈10^−12^, 95% Cl_1_ [0.06, 0.10], W_2_=2387.0, p_2_≈10^−15^, 95% Cl_2_ [0.09, 0.12], W_3_=2408.5, p_3_≈10^−15^, 95% Cl_3_ [0.07, 0.11], W_4_=2258.5, p_4_≈10^−12^, 95% Cl_4_ [0.05, 0.08]* and 12-dendrite vs. 3-dendrite: Wilcoxon test, *W_1–4_=2500.0, p_1–4_≈10^−16^, 95% Cl_1_ [0.24, 0.28], 95% Cl_2_ [0.22, 0.26], 95% Cl_3_ [0.20, 0.24], 95% Cl_4_ [0.16, 0.19]*). Both sparsity and pattern separation efficiency were highest in the 12-dendrite, followed by the 6-dendrite and the 3-dendrite models. The same was seen when using the rate-based distance metric to evaluate sparsity and pattern separation (Figure 6-figure supplement 1). In all cases, differences were more pronounced in the growth compared to the pruning case, in line with the findings of Figures 4–5.

**Figure 6.**
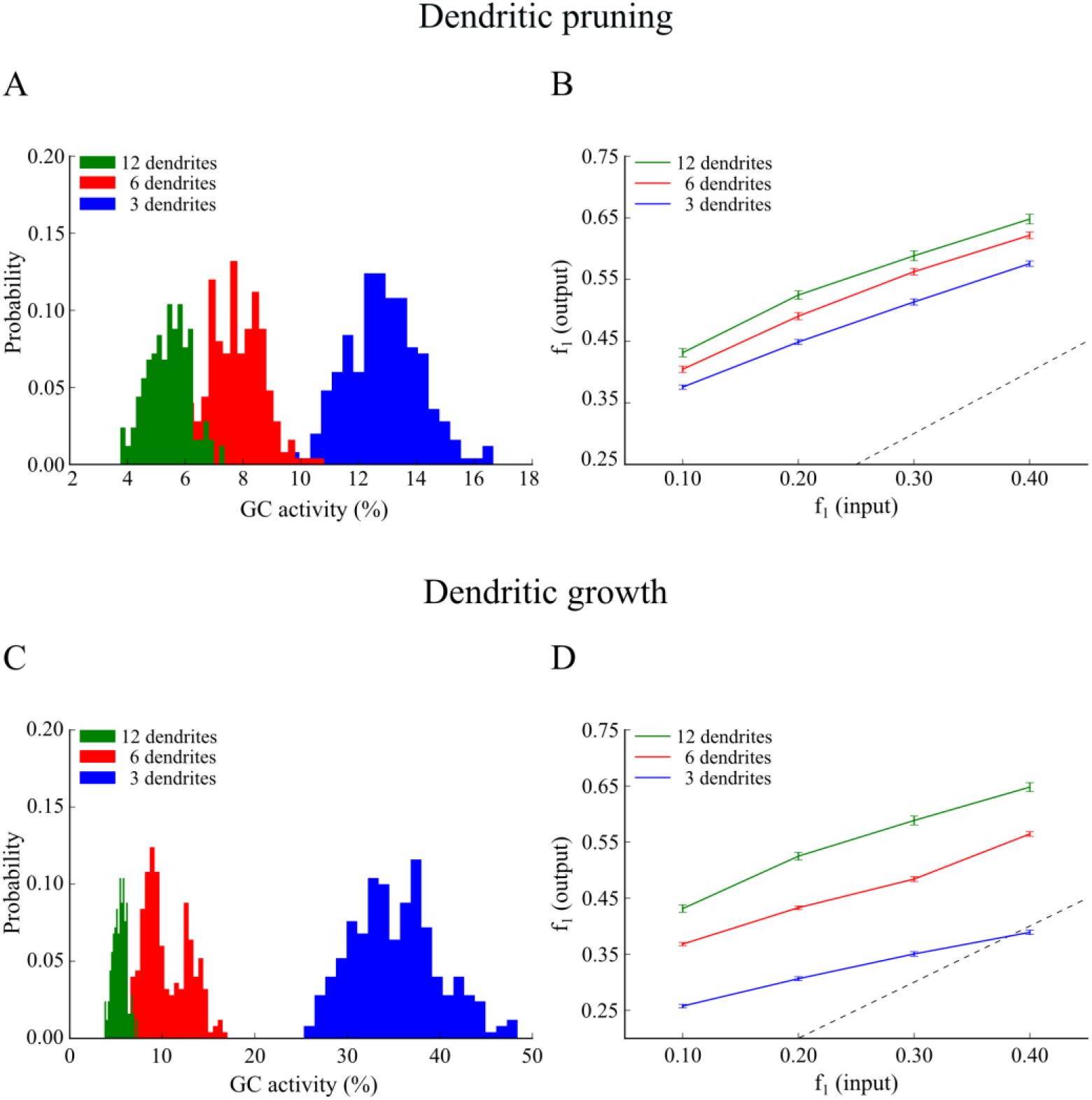
Effect of GC dendritic pruning (top panel) and growth (bottom panel) on pattern separation when the input resistance (R_in_) is the same across models. To match *R_in_*, the leak conductance (g_leak_) was increased by a factor of 1.695 and 1.230, in the 3- and 6-dendrite models respectively. **A**. Corresponding probability density functions of GC activity for the three GC models in response to a single input pattern at 40 Hz. GC activity distributions become more similar but remain inversely analogous to the number of dendrites: as the number of dendrites increase, the GC population becomes sparser. The respective mean firing rates of GCs are shown in Table 3. **B**. Input / Output population distances (*f*_1_), for the 3-dendrite (blue), 6-dendrite (red) and 12-dendrite (green) GC models in response to the presentation of two overlapping input patterns at 40 Hz as depicted in Figure 1B. The dashed line denotes the limit above which the model performs pattern separation. Performance improves with the number of dendrites. **C**. The same as in **A**. for the growth experiment. While distributions move closer to one another, the inverse relationship between dendritic number and mean sparsity is preserved. **D**. Same as in **B**. for the growth experiment. Pattern separation efficiency still correlates with the number of dendrites. Figure 6-figure supplement 1 depicts the corresponding pattern separation efficiency based on ‘rate distance’.

Similar results were obtained when correcting the input resistance by increasing the dimensions of the somatic compartment (Supplementary file 1B). For the pruning models, the soma increased by a factor of *1.278* and *1*.527 in the 6- and 3-dendrite models, respectively while for the growth models, these factors were *1.513* and *1.746*, respectively. As shown in Figure 7 (A-B), for the pruning case, differences in sparsity (12-dendrite vs. 6-dendrite: Kolmogorov-Smirnov test, *D=0.820, p≈10^−16^* and 12-dendrite vs. 3-dendrite, Kolmogorov-Smirnov test, *D=1.000, p≈10^−16^*) and pattern separation efficiency (12-dendrite vs. 6-dendrite: Wilcoxon test, *W_1_=1420.5, p_1_=0.2412, 95% Cl_1_ [−0.01, 0.03], W_2_=1588.0, p_2_=0.0020, 95% Cl_2_ [0.01, 0.04], W_3_=1713.5, p_3_=0.0014, 95% Cl_3_ [0.02, 0.05], W_4_=1600.5, p_4_=0.0158, 95% Cl_1_ [0.01, 0.04]* and 12-dendrite vs. 3-dendrite: Wilcoxon test, *W_1_=1914.0, p_1_≈10^−6^, 95% Cl_1_ [0.03, 0.07], W_2_=1878.0, p_2_≈10^−5^, 95% Cl_2_ [0.02, 0.06], W_3_=1979.5, p_3_≈10^−7^, 95% Cl_3_ [0.03, 0.07], W_4_=1891.5, p_4_≈10^−5^, 95% Cl_4_ [0.02, 0.05])* among corrected models decreased significantly but were not eliminated. Similar findings were seen in the growth case, both for the sparsity (12-dendrite vs. 6-dendrite: Kolmogorov-Smirnov test, *D=1.000, p≈10^−16^* and 12-dendrite vs. 3-dendrite: Kolmogorov-Smirnov test, *D=1.000, p≈10^−16^)* and pattern separation efficiency (12-dendrite vs. 6-dendrite: Wilcoxon test, *W_1_=1965.0, p_1_≈10^−13^, 95% Cl_1_ [0.04, 0.08], W_2_=1961.5, p_2_≈10^−6^, 95% Cl_2_ [0.03, 0.07], W_3_=2112.5, p_3_≈10^−9^, 95% Cl_3_ [0.04, 0.08], W_4_=1919.5, p_4_≈10^−6^, 95% Cl_4_ [0.02, 0.06]* and 12-dendrite vs. 3-dendrite: Wilcoxon test, *W_1_=2465.0, p_1_≈10^−16^ 95% Cl_1_ [0.12, 0.15], W_2_=2463.0, p_2_≈10^−16^, 95% Cl_2_ [0.10, 0.14], W_3_=2470.0, p_3_≈10^−16^, 95% Cl_3_ [0.09, 0.12], W_4_=2388.0, p_4_≈10^−15^95% Cl_4_ [0.07, 0.10])*. Both sparsity and pattern separation efficiency remained highest in the 12-dendrite, followed by the 6-dendrite and the 3-dendrite models. Similar findings were obtained when using the rate-based distance metric to assess pattern separation (Figure 7-figure supplement 1). Note that correcting the input resistance via increasing the somatic compartment is more effective than increasing the leak conductance, as both C_m_ and g_leak_ increase proportionally, keeping the membrane time constant identical across model cells.

**Figure 7.**
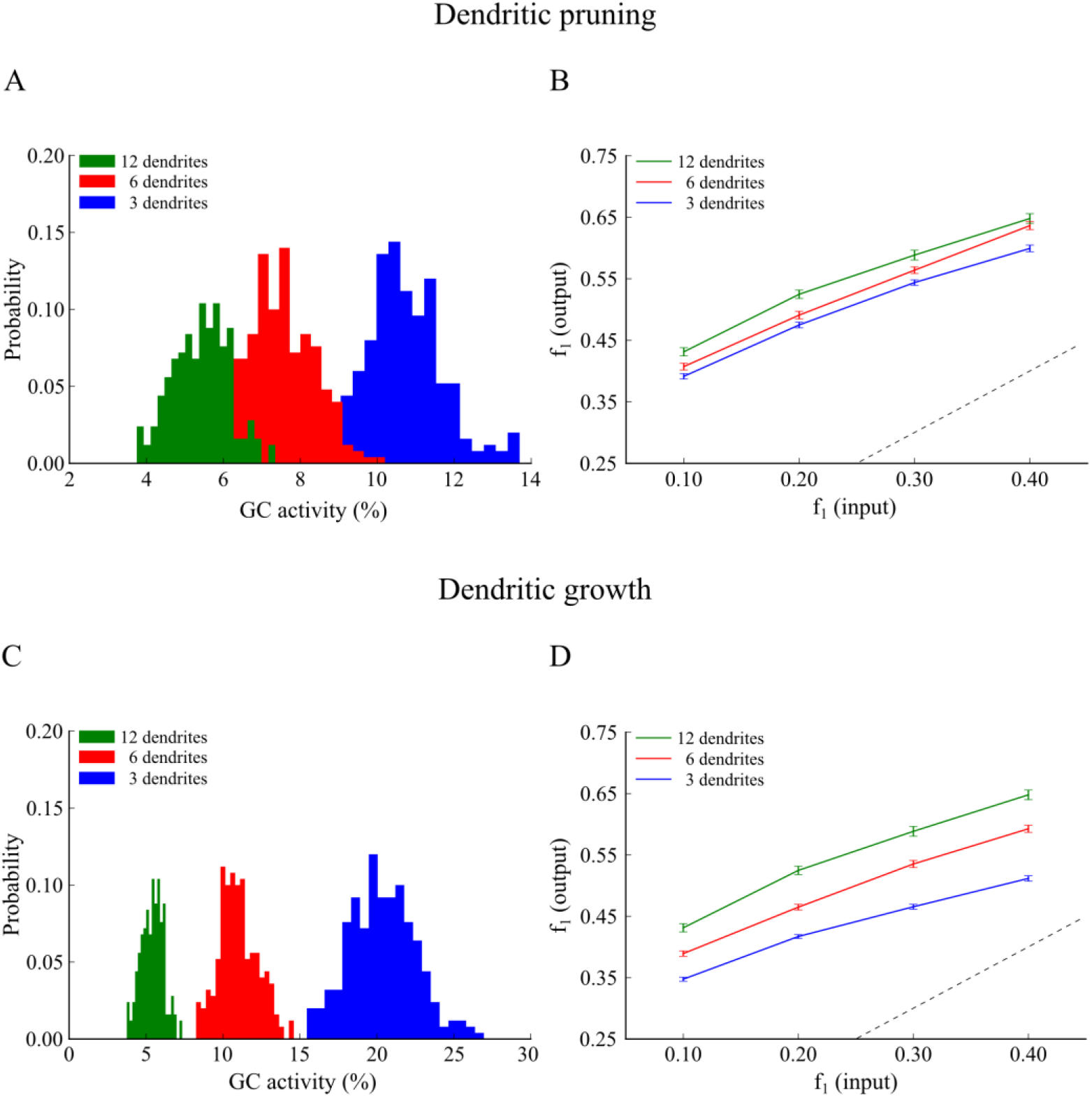
Effect of GC dendritic pruning (top panel) and growth (bottom panel) on pattern separation when matching the input resistance (R_in_) via increasing the somatic size. The neuronal soma of the 3- and 6-dendrite models was increased by a factor of 1.527 and 1.278, respectively. **A**. Corresponding probability density functions in response to a single input pattern at 40 Hz. While distributions move closer, the mean activity remains inversely analogous to the number of dendrites. The respective mean firing rates of GCs are shown in Table 3. **B**. Input / Output population distances (*f*_1_), for the 3-dendrite (blue), 6-dendrite (red) and 12-dendrite (green) GC models in response to the presentation of two overlapping input patterns at 40 Hz as depicted in Figure 1B. The dashed line denotes the limit above which the model performs pattern separation. Performance becomes similar yet statistically different among the three corrected models and remains analogous to the number of dendrites. **C**. Same as in **A**. for the growth experiment. The inverse relationship between dendritic number and mean sparsity is preserved. **D**. Same as in **B**. for the growth experiment. Pattern separation efficiency remains different and analogous to the number of dendrites. Figure 7-figure supplement 1 depicts the corresponding pattern separation efficiency based on ‘rate distance’.

Overall, these simulations suggest that while the input resistance is a key determinant of GC neuron activity, it does not fully explain the differences in sparsity and pattern separation efficiency among models with 3, 6 or 12 GC dendrites.

#### Sparsity is the key determinant of pattern separation efficiency

The question that arises naturally from the above findings is whether further manipulations of the leak conductance and/or somatic dimensions could match sparsity across all models? Moreover, would matching sparsity result in identical pattern separation efficiency, thus making sparsity the key determinant of pattern separation (O’Reilly and McClelland, 1994; Johnston et al., 2015)? To answer these questions we explored the effects of manipulating intrinsic (g_leak_ and somatic dimensions) as well as extrinsic (synaptic weight) mechanisms in the 3-, 6- and 12-dendrite GC models. Due to the consistent nature of the previous results, the remaining simulations were performed using the pruning experiment configuration to generate the 3- and 6-dendrite GC models and the population-based metric to assess pattern separation. Their corresponding properties are shown in Supplementary file 1C and the corrected dendritic integration profiles in Figure 2-figure supplement 5.

As shown in Figure 8Ai, increasing g_leak_ by a factor of *1.58* and *2.48*, in the 6- and 3-dendrite GC models respectively, eliminated the differences in sparsity distributions compared to the control (12-dendrite vs. 6-dendrite: Kolmogorov-Smirnov test, *D=0.096, p=0.1995* and 12-dendrite vs. 3-dendrite: Kolmogorov-Smirnov test, *D=0.100, p=0.1641*). The same was observed with respect to pattern separation efficiency (Figure 8Aii). All three models exhibited identical performance across all difficulty levels (12-dendrite vs. 6-dendrite: Wilcoxon test, *W_1_=1148.5, p_1_=0.4863, 95% Cl_1_ [−0.03, 0.01], W_2_=1263.5, p_2_=0.9286, 95% Cl_2_ [−0.02, 0.02], W_3_=1337.5, p_3_=0.5486, 95% Cl_3_ [−0.01, 0.02], W_4_=1192.0, p_4_=0.6918, 95% Cl_4_ [−0.02, 0.02]*, 12-dendrite vs. 3-dendrite: Wilcoxon test, *W_1_=1195.5, p_1_=0.7097, 95% Cl_1_ [−0.03, 0.01], W_2_=1067.5, p_2_=0.2096, 95% Cl_2_ [−0.04, 0.01], W_3_=1326.5, p_3_=0.6003, 95% Cl_3_ [−0.01, 0.02], W_4_=1246.5, p_4_=0.9835, 95% Cl_4_ [−0.02, 0.02])*. Similarly, increasing the diameter and length of the somatic compartment in the 6- and 3-dendrite models by *1.480* and *1.870*, respectively (Figure 8Bi) resulted in similar sparsity (12-dendrite vs. 6-dendrite: Kolmogorov-Smirnov test, *D=0.108, p=0.1083* and 12-dendrite vs. 3-dendrite: Kolmogorov-Smirnov test, *D=0.040, p=0.9883)*. Similarly, pattern separation efficiency was identical across all models and difficulty levels (Figure 8Bii), (12-dendrite vs. 6-dendrite: Wilcoxon test, *W_1_=1092.0, p_1_=0.2776, 95% Cl_1_ [−0.04, 0.01], W_2_=1106.5, p_2_=0.3242, 95% Cl_2_ [−0.03, 0.01], W_3_=1350.5, p_3_=0.4906, 95% Cl_3_ [−0.01, 0.03], W_4_=1378.0, p_4_=0.3794, 95% Cl_4_ [−0.01, 0.03]* and 12-dendrite vs. 3-dendrite: Wilcoxon test, *W_1_=1144.0, p_1_=0.4670, 95% Cl_1_ [−0.03, 0.01], W_2_=937.5, p_2_=0.0315, 95% Cl_2_ [−0.04, 0.00], W_3_=1281.0, p_3_=0.8335, 95% Cl_3_ [−0.02, 0.02], W_4_=1117.5, p_4_=0.3628, 95% Cl_4_ [−0.03, 0.01]*). It should be noted that the abovementioned sizes are not realistic for the somata of GC neurons. Nevertheless, these findings highlight the key role of sparsity in controlling pattern separation efficiency, irrespectively of the number of GC dendrites. Moreover, these simulations predict that intrinsic mechanisms of GC neurons could potentially be used to correct for morphological alterations in order to control sparsity levels.

**Figure 8.**
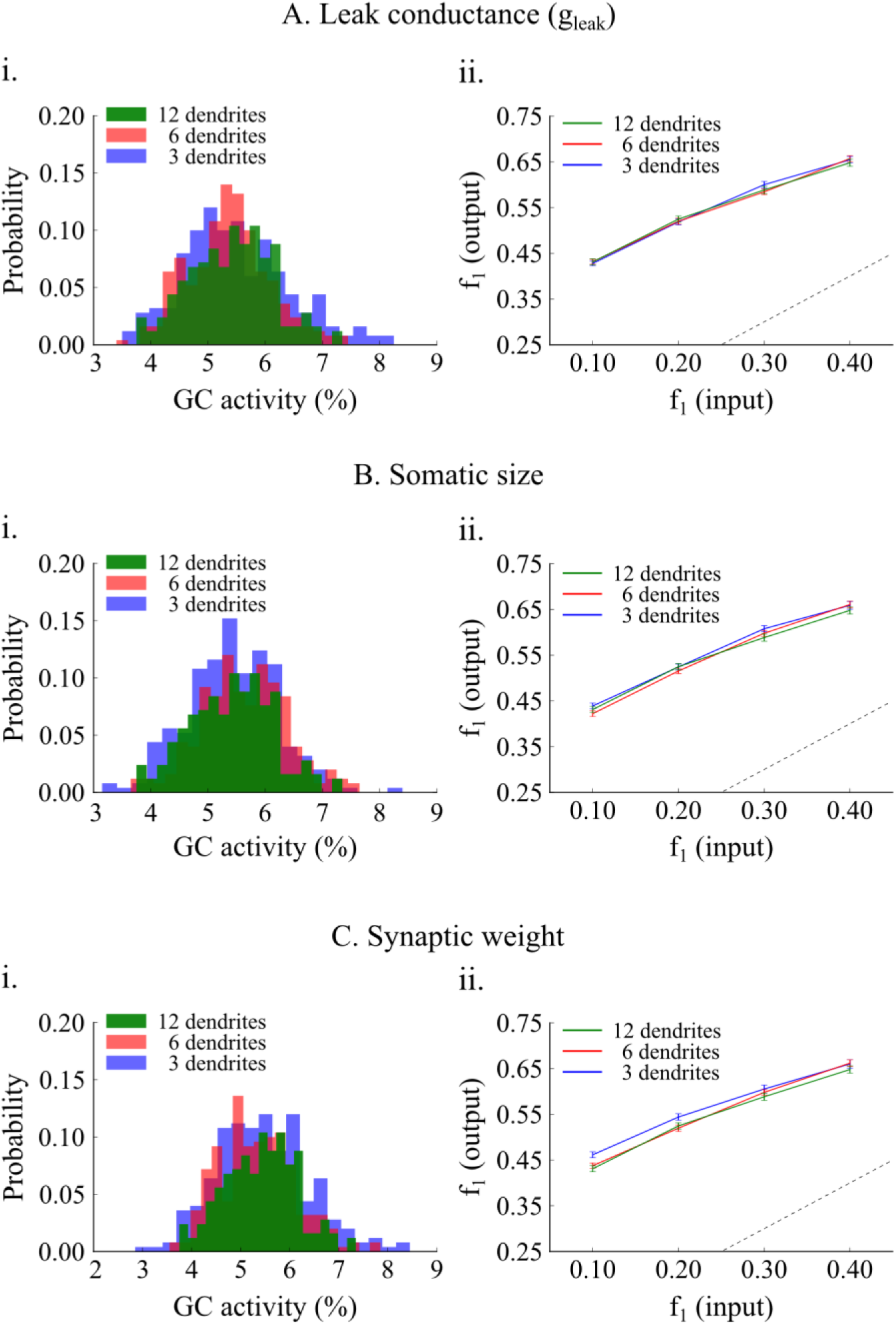
Effect of matching sparsity on pattern separation efficiency. **A**. The “leak” conductance, g_leak_, of the 6- and 3-dendrite GC models increases by a factor of 1.58 and 2.48, respectively. **A.i** Resulting GC activity distributions are not statistically different. **A.ii** Networks with corrected GC models have identical performance. Differences in pattern separation using the population metric are not statistically significant. **B**. GC activity distributions are matched by increasing the soma diameter and length of the 6- and 3-dendrite GC models by 1.48 and 1.87, respectively. **B.ii** The network with corrected 3-dendrite GC models has a slightly smaller pattern separation efficiency compared to the other two probably because its activity distribution is wider. Generally the three models have very similar pattern separation performance. **C**. GC activity distributions are matched by decreasing the EC ➔ GC synaptic weight from 1.00, to 0.75 and 0.56 in the 6-, and 3-dendrite models, respectively. Pattern separation efficiency is statistically the same across all corrected GC models.

We next investigated whether sparsity levels can also be controlled via extrinsic (network) rather than intrinsic mechanisms such as changes in the synaptic weights between EC and GC neurons, namely by decreasing the strength of the input to the DG network. Specifically, the synaptic weight of the EC ➔ GC synapses in the 6- and 3-dendrite models was set to *0.75* and *0.56*, respectively (the control value in the 12-dendrite model was *1.00*). Again, as shown in Figure 8Ci, differences in sparsity were eliminated among the three dendrite models (12-dendrite vs. 6-dendrite: Kolmogorov-Smirnov test, *D=0.092, p = 0.2406* and 12-dendrite vs. 3-dendrite: Kolmogorov-Smirnov test, *D=0.092, p=0.2406*). Finally, pattern separation efficiency was nearly identical cross all models and difficulty levels (Figure 8Cii), with a tiny difference in the 3-dendrite model for patterns overlapping by 90% (12-dendrite vs. 6-dendrite: Wilcoxon test, *W_1_=1085.5, p_1_=0.2582, 95% Cl_1_ [−0.04, 0.01], W_2_=1072.5, p_2_=0.2224, 95% Cl_2_ [−0.03, 0.01], W_3_=1316.5, p_3_=0.6491, 95% Cl_3_ [−0.02, 0.02], W_4_=1122.0, p_4_=0.379495% Cl_4_ [−0.03, 0.01]* and 12-dendrite vs. 6-dendrite: Wilcoxon test, *W_1_=1117.5, p_1_=0.3628, 95% Cl_1_ [−0.04,0.01], W_2_=1052.5, p_2_=0.1744, 95% Cl_2_ [−0.04, 0.01], W_3_=966.0, p_3_=0.0507, 95% Cl_3_ [−0.01, 0.01], W_4_=773.0, p_4_=0.0010, 95% Cl_4_ [−0.05, −0.01)]*).

In sum, these simulations predict that sparsity is the key determinant of pattern separation efficiency. Sparsity of the GC neuronal population can be controlled via multiple mechanisms, including the growth of dendrites. Pruning of dendrites can be compensated by the growth of very large somata, a large increase in the leak channel conductance or a significant decrease in the strength of the EC input via synaptic weight changes.

### What have we learnt from the model?

The above simulations suggest that dendrites contribute to pattern separation via enhancing sparsity, yet they are not essential for this task. Different mechanisms, both intrinsic and extrinsic, can achieve the same effects. What is not entirely clear is whether the relationship between sparsity and pattern separation efficiency is identical for all three GC models across different sparsity levels. To test this we challenged the three models (deduced from the pruning experiment) with the task of separating pairs of inputs that overlap by 80%, while varying the synaptic weight of the EC ➔ GC connections so as to induce different levels of sparsity. Only corrected models (namely exhibiting the same level of sparsity) were compared. As shown in Figure 9A, for any given sparsity level, pattern separation efficiency was almost identical for all three dendrite models (Wilcoxon test, *p>0.05*). These simulations suggest that indeed, when it comes to pattern separation, the key determinant is the level of GC population sparsity. Having multiple dendrites simply helps achieving high levels of sparsity because the probability of generating somatic spikes when inputs are distributed across many (rather than few) dendrites is significantly smaller.

Figure 9B summarizes the predictions made by our DG model, whereby sparsity plays a central role in pattern separation. We already knew from previous studies that inhibition is one of the key mechanisms mediating sparsity and in turn pattern separation (Myers and Scharfman, 2009, 2011; Jinde et al., 2012). What we propose here is that the presence of dendrites comes with a bonus in the DG network: it provides another mechanism for increasing sparsity and as such has a key role in pattern separation. While dendrites appear to be sufficient, they are not necessary for achieving high sparsity levels. Both intrinsic and network mechanisms can be used to achieve the same effects. In this study we identified some of these mechanisms but it is likely that there are many others and/or their combinations can also achieve similar results.

**Figure 9.**
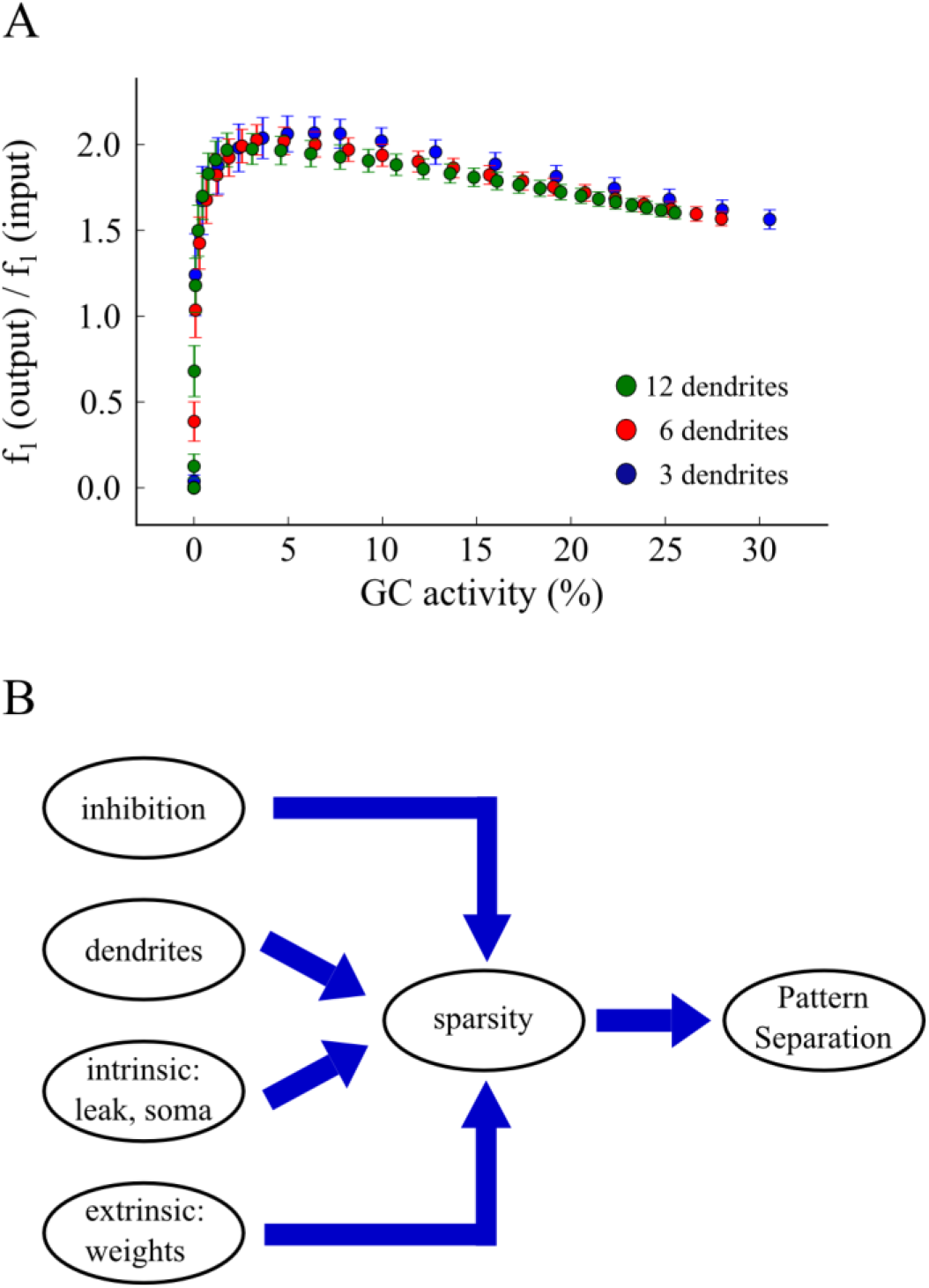
**A**. The ratio of *f*_1(output)_ to *f*_1(input)_ for all three models as function of GC activity. Each dot represents an average of 20 trials for a specific synaptic weight value. We used values in the range of 0.0 to 25.0. For any given sparsity level, pattern separation efficiency is identical across all corrected GC models. **B**. Schematic representation of our proposed working hypothesis based on model predictions: sparsity is the key determinant of pattern separation. Sparsity is in turn controlled by a number of mechanisms, including inhibition, the presence of dendrites as well as various intrinsic and extrinsic mechanisms.

## Discussion

### Summary of the results

#### The model

The goal of this study was to reveal whether and how dendrites of principal (GC) cells can mediate pattern separation in the DG via the use of a computational approach. Towards this goal, we introduced a novel network model of the DG that includes the four major neuronal cell types found in this area, namely Granule Cells, Mossy Cells, Basket Cells, and HIPP cells. GCs were modeled as two stage integrators via the addition of dendritic branches whose properties are loosely constrained by electrophysiological and anatomical data. The rest of the neuronal types were simulated as exponential leaky integrate-and-fire neurons with adaptation. The proposed hybrid model serves as a bridge between simplified point neuron network models (Myers and Scharfman, 2009) and more detailed biophysical models (Santhakumar et al., 2005) of the DG. As such, it provides a biologically relevant and computationally efficient tool for the in depth exploration of different factors that may contribute to pattern separation, going beyond the scope of this particular study. The selective use of dendritic compartments only in GCs keeps the model complexity low while at the same time allowing the dissection of some basic GC dendritic mechanisms in pattern separation. To our knowledge this is the first DG network model of its kind.

#### Model predictions

Inhibition is known to control neuronal activity by increasing sparsity, as the number of active neurons is smaller for a given stimulus (Jung and McNaughton, 1993) therefore, enhancing pattern separation (Aimone et al., 2011). In the presented model, inhibition is provided to the network both directly via BCs (perisomatic inhibition) and HIPP cells (dendritic inhibition) and indirectly through the MC circuitry. We find that MC loss, while increasing GC cell activity in line with the experimental data of (Ratzliff et al., 2002), does not lead to hyper-excitability yet we do predict a measurable deficit on pattern separation.

To our knowledge, our model is the first to predict a role of GC dendrites in pattern separation. This role is an indirect one and results from the inherent increase in sparsity of the GC cells that is endowed by the presence of dendrites. Specifically, we show that the number of GC dendrites correlates positively with pattern separation efficiency due to the higher sparsity levels provided by having multiple dendrites. In our control model, higher sparsity arises from the requirement of having at least two dendrites simultaneously active in order to fire a GC model neuron. This emerged from the calibration of GC properties against experimental data, thus is considered biologically relevant. As a result, GCs with large numbers of dendrites have a lower probability of activation given a fixed number of afferents, therefore increased network sparsity. We also predict that under conditions of dendritic pruning and/or early in the growth stages of GCs, high sparsity can be achieved with alternative mechanisms, both intrinsic (e.g., leak conductance, somatic dimensions) and extrinsic (e.g., synaptic weights) making dendrites a sufficient but not necessary condition for high pattern separation efficiency. These results support the hypothesis that sparsity in GC activity improves pattern separation (O’Reilly and McClelland, 1994) and provide a list of alternative mechanisms for controlling sparsity in the DG.

### Implications for pathology

As part of the hippocampus, the DG is long hypothesized to play a key role in associative memories, and especially when those are related with events (Morris, 2006). Furthermore, the hippocampal DG has been implicated as the subregion most sensitive to the effects of advancing age (Small et al., 2004). While the CA1 subregion is directly associated with Alzheimer’s Disease (AD) due to cell loss, as demonstrated in humans (West et al., 2006), DG alterations have also been reported in patients with the aforementioned disease (Scheff and Price, 2003), including changes in granule cell dendrites (Einstein et al., 1994).

Specifically, the dendrites of GCs in AD patients appear shorter, with fewer branches and fewer spines than those of matched controls. Moreover, in AD patients the dendrites of granular cells were reported to lose approximately 50% of their spines (Einstein et al., 1994). Our simulations show that the most important of the above-listed observations with respect to pattern separation performed by the DG would be the shortening of dendritic branches via the loss of branch points rather than the loss of side branches while mentioning the same dendritic length. Total dendritic length of GCs was previously linked to AD, which in turn is aligned with the evidence that patients with AD, who have extensive hippocampal and parahippocampal damage, lost their ability to encode information in distinct, orthogonal representations (Ally et al., 2013).

The DG is also associated with epileptogenesis in temporal lobe epilepsy (TLE) and hence, many computational models are used to investigate the effect of GC alterations in epilepsy (Tejada and Roque, 2014; Faghihi and Moustafa, 2015). Moreover, hilar cell loss has been reported in animal models after concussive head injury and also under TLE (Mathern et al., 1995). It remains unclear however, which hilar neurons are lost in animal models of TLE. As a result, there are currently three theories for TLE: a) the “dormant basket cell” hypothesis according to which the hyper-excitability in GC population is due to the loss of MCs which normally excite BCs which in turn provide inhibition to GCs. b) The “irritable mossy cell” hypothesis according to which surviving MCs hyper-excite GCs by sending uncontrolled excitation, and c) the MC loss-induced sprouting hypothesis (mossy fiber sprouting) (Ratzliff et al., 2002). We show that under conditions of MC loss, GCs exhibit increased activity (but not hyper-excitability) which should lead to pattern separation deficits. While our results are in agreement with the findings of (Ratzliff et al., 2002), more experiments need to be performed to revolve this debatable issue.

### Simplifications of the model, and future directions

Several simplifications were made in modeling the individual cells and in implementing the DG network. First, we used simple point neurons in order to simulate the neuronal cells of DG. Although these models could capture the average spiking properties of a given neuron, it remains unclear how the geometrical characteristics of those neurons could affect their behavior. Another simplification concerns the effects of synaptic failure rates and receptor desensitization (Harney and Jones, 2002) in the DG, which were not included in the model.

An important aspect of DG function is the long-term synaptic plasticity, by which the connections from PP to GCs are modified. Previous DG models used a form of Hebbian learning that incorporates features of long-term potentiation and depression (Rolls, 2007). However, such a function is most likely to be relevant when stimuli are presented repetitively. On the contrary, the current model is used to distinguish patterns presented in single instances and accordingly, plasticity is not considered. Future work may address such issues along with including other interneuronal populations such as the Molecular layer Perforant Path-associated (MOPP) and the Hilar Commissural-Associational pathway related (HICAP) cells, especially when more data on their intrinsic and connectivity properties become known.

Moreover, GCs are among few cells that undergo neurogenesis in adulthood (Eriksson, 2003; Aimone et al., 2010). In a recent study by Nakashiba et al. (2012), the role of young GCs in pattern separation was investigated and it was concluded that new neurons are required for the discrimination of similar inputs. Since the presented model is used to examine specific alterations of GCs and their effect on pattern separation, neurogenesis is not incorporated but would be considered in the future. Overall, the abovementioned simplifications are unlikely to have a major effect on the basic conclusions about the contribution of morphological alterations of GC dendrites to pattern separation.

## Materials and Methods

The model was developed mainly based on the structure and connectivity features described by Myers and Scharfman (2009), and incorporates the four major dentate cell types. These are the GCs, MCs, BCs and HIPP cells. As the Hilar Commissural-Associational Pathway (HICAP) cells are relatively rare and poorly understood (Sik et al., 1997), they are not included in the model. All simulations were performed using the BRIAN (BRIAN v1.4) network simulator (Goodman and Brette, 2009; Brette and Goodman, 2011) running on a High-Performance Computing Cluster (HPCC) with 312 cores under 64-bit CentOS Linux.

### Model neurons

The four types of DG neurons were modeled as simplified phenomenological neurons of the integrate-and-fire (I&F) type (Izhikevich, 2003; Burkitt, 2006), with no internal geometry (“point neurons”). The GCs incorporated dendrites in order to study their role in pattern separation; however the MCs, BCs and HIPP cells were simulated as simple somatic compartments.

### Modeling BC, MC and HIPP cells

Specifically, an adaptive exponential I&F model (aEIF) (Brette and Gerstner, 2005) was used to model MCs, BCs and HIPP cells. The model is mathematically described by the following differential equations (Equation 1, 2):

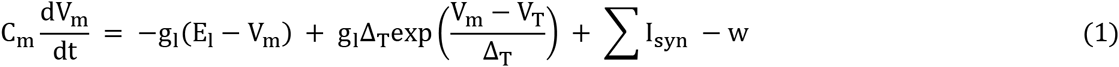

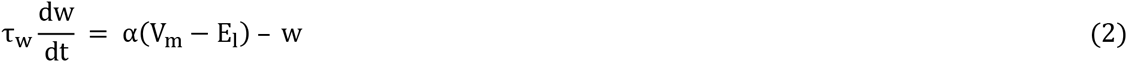

where C_m_ is the membrane capacitance, V_m_ the membrane voltage, g_l_ the “leak” conductance, E_l_ the “leak” reversal potential (i.e., the resting potential), I_syn_ the synaptic current flow onto the neuron, w the adaptation variable, Δ_T_ the slope factor, V_T_ the effective threshold potential, α the adaptive coupling parameter, and τ_w_ is the adaptation time constant.

The exponential nonlinearity describes the spike action potential and its upswing. In the mathematical interpretation of the model a spike occurs at time t_spike_ when the membrane voltage reaches a finite limit value, and thereafter the downswing of the action potential is described by a reset fixed value V_reset_, as follows:

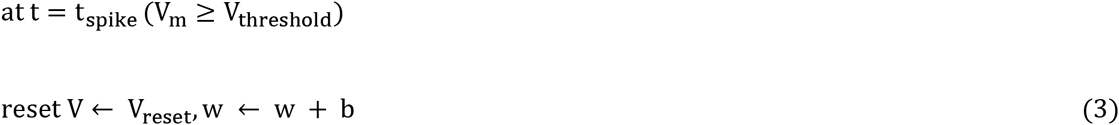

where V_threshold_ is the voltage threshold above which the neuron fires a spike, and b is the spike triggered adaptation parameter. For all neuron types the effective threshold is equal to the voltage threshold (see Table 4 for model parameters).

**Table 3.**
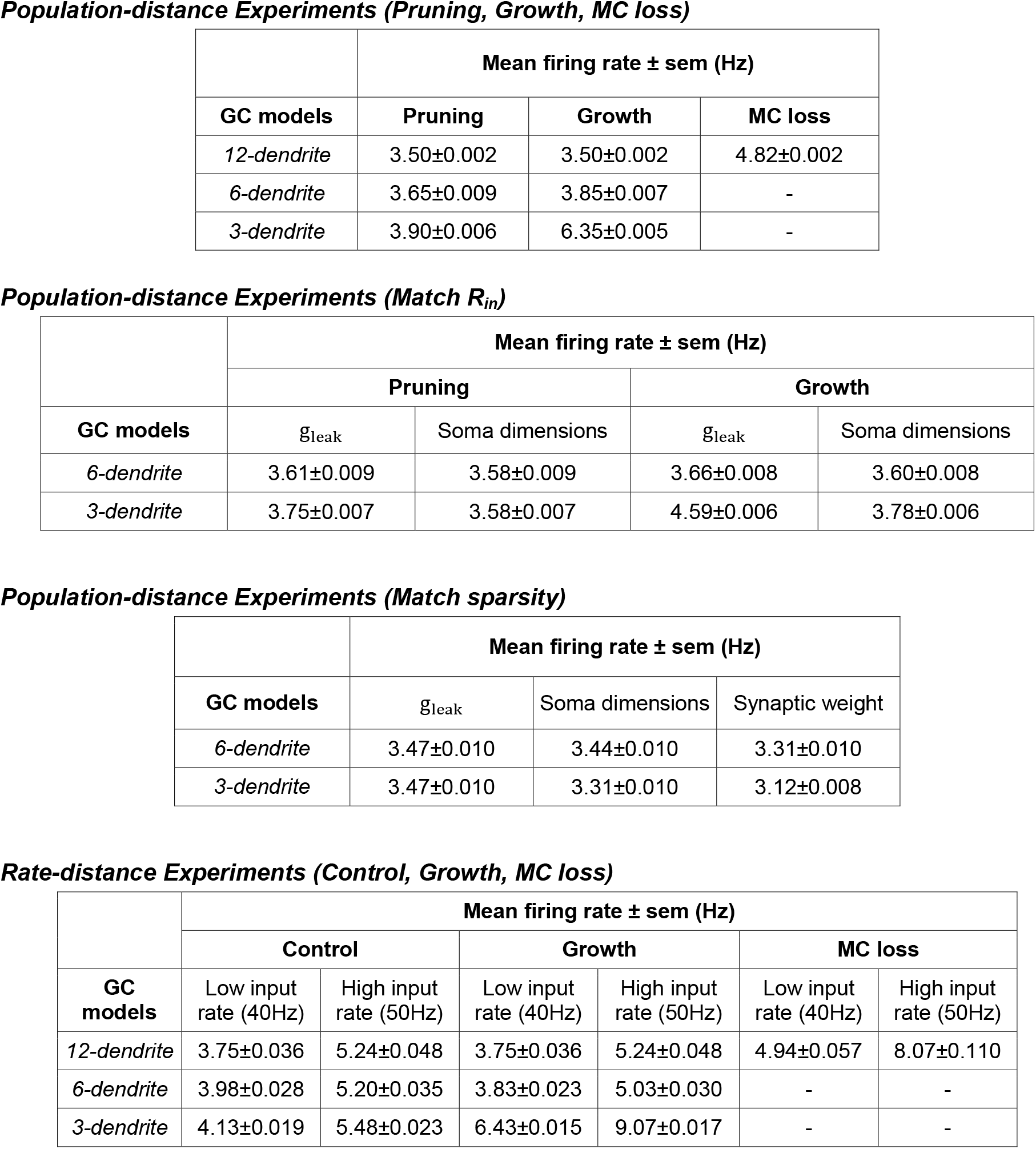
Mean firing frequencies of GCs in response to the various input patterns used in Figures 3–8

**Table 4.** Model parameters for all neuronal types

| Model Parameter | Granule cells |  | Mossy cells | Basket cells | HIPP cells |
| --- | --- | --- | --- | --- | --- |
|  | Soma | dendrites |  |  |  |
| $E_l$ (mV) Resting potential | -87 | -82 | -64 | -52 | -59 |
| $g_l$ (nS) "Leak" conductance | 0.00003<sup>a</sup> | 0.00001<sup>a</sup> | 4.53 | 18.054 | 1.930 |
| $C_m$ (nF) Membrane capacitance | 1.0<sup>b</sup> | 2.5<sup>b</sup> | 0.621 | 0.1793 | 0.0584 |
| $V_{reset}$ (mV) Reset voltage | -74 | | -49 | -45 | -56 |
| $V_T = V_{thr}$ (mV) Threshold voltage | -56 | | -42 | -39 | -50 |
| $\Delta_T$ (mV) Slope factor | - | | 2 | 2 | 2 |
| $\alpha$ (nS) Adaptation coupling parameter | 2.0 | | 2 | 0.1 | 0.82 |
| $\tau_w$ (ms) Adaptation time constant | 45 | | 180 | 100 | 93 |
| $b$ (nS) Spike triggered adaptation | 0.0450 | | 0.0829 | 0.0205 | 0.015 |

<sup>a</sup>For the GC model the "leak" conductance is given in Siemens/cm<sup>2</sup>
<sup>b</sup>For the GC model the membrane capacitance is given in uF/cm<sup>2</sup>

### Modeling principal neurons, GC

In order to investigate the role of GC dendrites in pattern separation, an extended point neuron was implemented. The GC model consisted of a leaky Integrate-and-Fire somatic compartment connected to a variable number of dendritic compartments whose morphology relies on anatomical data (see Table 5 for structure characteristics). Furthermore, an adaptation parameter w was used, only in the somatic compartment, to reproduce spike frequency adaptation reported in these neurons. The equation that describes the membrane, somatic and dendritic, potential of GC model cells is as follows:

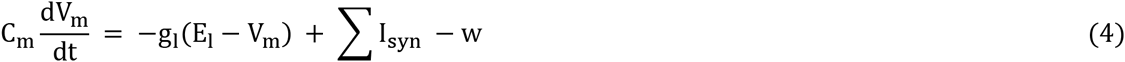

The adaptation parameter (w) was set to zero for the dendritic compartments. There is no evidence for dendritic spikes in GCs (Krueppel et al., 2011), thus the spike mechanism was only applied in the somatic equation.

The DG is divided into three distinct layers (Figure 1A), the molecular, granular, and polymorphic (hilus) (Amaral et al., 2007). The GC dendrites extend in the molecular layer (Amaral et al., 2007), which is further divided into the inner, middle, and outer molecular layers, and therefore dendritic compartments are discretized accordingly. Table 5 lists the morphological characteristics of the GC model. According to anatomical data, (Claiborne et al., 1990) GCs have 10–15 dendrites; thus, the control GC model includes 12 dendrites and its physiological responses are validated against experimental data (see Table 4 for GC model parameters).

In order to investigate whether the number of GC dendrites affects pattern separation, we used two different approaches; dendritic pruning and growth. First, two more GC models were implemented which differ only in their number of dendrites (6 and 3), but the path length remained the same across these models. Secondly, two GC models were implemented which differ both in their dendritic number and their path length. The morphological differences among the three models are shown in Table 5. The membrane capacitance of the dendritic compartments was increased compared to the somatic one in order to account for spines reported in GC dendrites (Aradi and Holmes, 1999).

**Table 5.** Morphological structure of GC models

| Structure of GC models | Control | Pruning |  | Growth |  |
| --- | --- | --- | --- | --- | --- |
|  | 12 dendrites | 6 dendrites | 3 dendrites | 6 dendrites | 3 dendrites |
| # of compartments |  |  |  |  |  |
| *total* | 21 | 15 | 9 | 9 | 3 |
| *proximal* | 3 | 3 | 3 | 3 | 3 |
| *medial* | 6 | 6 | 3 | 6 | - |
| *distal* | 12 | 6 | 3 | - | - |
| length per compartment (um) | 83 | 83 | 83 | 83 | 83 |
| total dendritic length (um) | 1743 | 1245 | 747 | 747 | 249 |
| diameter per compartment (um) |  |  |  |  |  |
| *proximal* | 1.0 | 1.0 | 1.0 | 1.0 | 1.0 |
| *medial* | 0.9 | 0.9 | 0.9 | 0.9 | - |
| *distal* | 0.8 | 0.8 | 0.8 | - | - |

The intrinsic model properties that were validated against experimental data are the input resistance (R_in_), the sag ratio, defined as the ratio between the exponentially extrapolated voltage to the steady-state voltage, and the membrane time constant (τ_m_). In line with experimental procedures (Lübke et al., 1998), we used 1-second somatic current injection to calculate the intrinsic properties. The input resistance is calculated by the equation R_in_ = ΔV_m_/I_injected_, where ΔV_m_ is the membrane response to current stimulation. Finally, the membrane time-constant is approximated by the formula τ_m_ = R_in_C_m_, which is a valid approximation for passive compartments. As experimental data were obtained in the presence of synaptic activity blockers, a somatic current injection at the model cell was used to replicate those conditions.

## Modeling Synapses

Since the DG network consists of both glutamatergic cells (GCs and MCs) and GABAergic interneurons, AMPA, NMDA and GABA synapses were included in the network model. Therefore, the total synaptic current (Equations 1 and 4) consisted of two components; the excitatory current through AMPA receptors (I_AMPA_) and NMDA receptors (I_NMDA_), and the inhibitory current via GABA_a_ receptors (I_GABA_). The majority of ligand-gated ion channels mediating synaptic transmission, such as AMPA and GABA receptors, display an approximately linear current-voltage relationship when they open. We modeled these channels as an ohmic conductance (g_syn_) multiplied by the driving force:

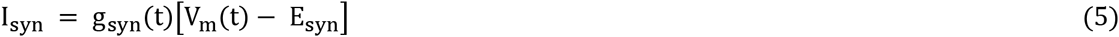

where E_syn_ is the AMPA and GABA reversal potential, respectively.

The NMDA receptor-mediated conductance depends on the postsynaptic voltage due to the gate blockage by a positively charged magnesium ion (Mg^2+^). The fraction of NMDA channels that are not blocked by Mg^2+^ can be fitted by a sigmoidal function (Jahr and Stevens, 1990):

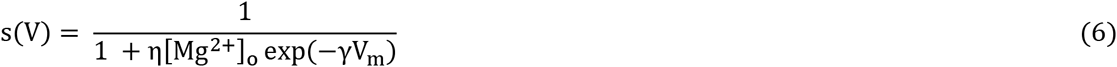

where η is the sensitivity of Mg unblock, γ the steepness of Mg unblock, and [Mg^2+^]_o_ is the outer magnesium (Mg) concentration. For NMDA receptors in MCs, BCs and HIPP cells we used η = 0.28 mM^−1^, [Mg^2+^] = 1mM, and γ = 0.072 mV^−1^. Instead, for GCs we tuned these parameters in to match the latest experimental data found in literature (Krueppel et al., 2011) with the corresponding values equal to η = 0.2 mM^−1^, [Mg^2+^] = 2mM, and γ = 0.04 mV^−1^. Consequently, the NMDA synaptic current is calculated by the following equation:

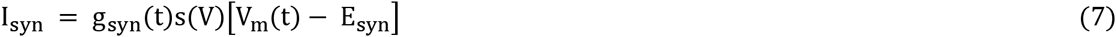

The ohmic conductance is simulated as a sum of two exponentials (Bartos et al., 2001), one term based on rising and the other on the decay phase of the postsynaptic potential. This function allows time constants to be set independently. We simulated such a function as a system of linear differential equations (Roth and Van Rossum, 2009):

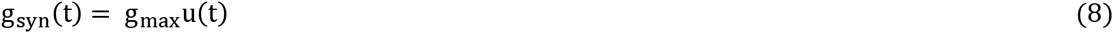

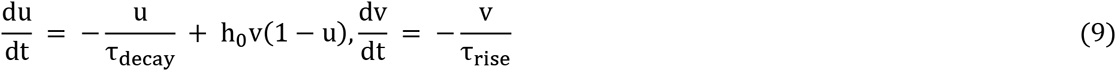

where τ_rise_ and τ_decay_ are the rise and decay constants respectively, h_0_ a scaling factor and u(t) is the function of two exponentials u(t) = exp(−t/τ_decay_) – exp(−t/τ_rise_), which is divided by its maximum amplitude. The scaling factor is set to 1 ms^−1^ for all AMPA and GABA receptors and all neuronal types. The NMDA scaling factor is set to 0.5 ms^−1^ apart from the synapses form on GCs where it is set to 2 ms^−1^. Because axons of neurons are not included in the model, a delay is used between pre- and postsynaptic transmission. The role of the delay is to account for both the synaptic transmission and the axonal conduction delay, and its value depends on the presynaptic and postsynaptic neuronal types. The peak conductance (g_max_), rise and decay time constants, and the delay of various network connections were estimated from experimental data (Kneisler and Dingledine, 1995; Geiger et al., 1997; Bartos et al., 2001; Schmidt-Hieber et al., 2007; Larimer and Strowbridge, 2008; Schmidt-Hieber and Bischofberger, 2010; Krueppel et al., 2011; Chiang et al., 2012) and are given in Table 6. Specifically, the GC peak conductance both for AMPA and NMDA, was validated against experimental data (Krueppel et al., 2011), where it is evidenced that a single synapse provokes a 0.6 mV somatic EPSP, and also the NMDA and AMPA peak current ratio is equal to 1.08. These values were reproduced in the GC model cells. The models also incorporate background activity, in order to simulate the experimental findings of spontaneous activity in DG. Accordingly, we used Poisson independent spike trains in order to reproduce the experimental data for MCs (2–4 Hz spontaneous activity) (Henze and Buzsáki, 2007) and for BCs (1–2 Hz spontaneous activity) (Kneisler and Dingledine, 1995). GCs infrequently generate spontaneously activity, even if inhibition is blocked (Lynch et al., 2000). Thus, we implemented noisy inputs in order to only evoke spontaneous EPSPs (0.05 Hz spontaneous activity).

**Table 6.**
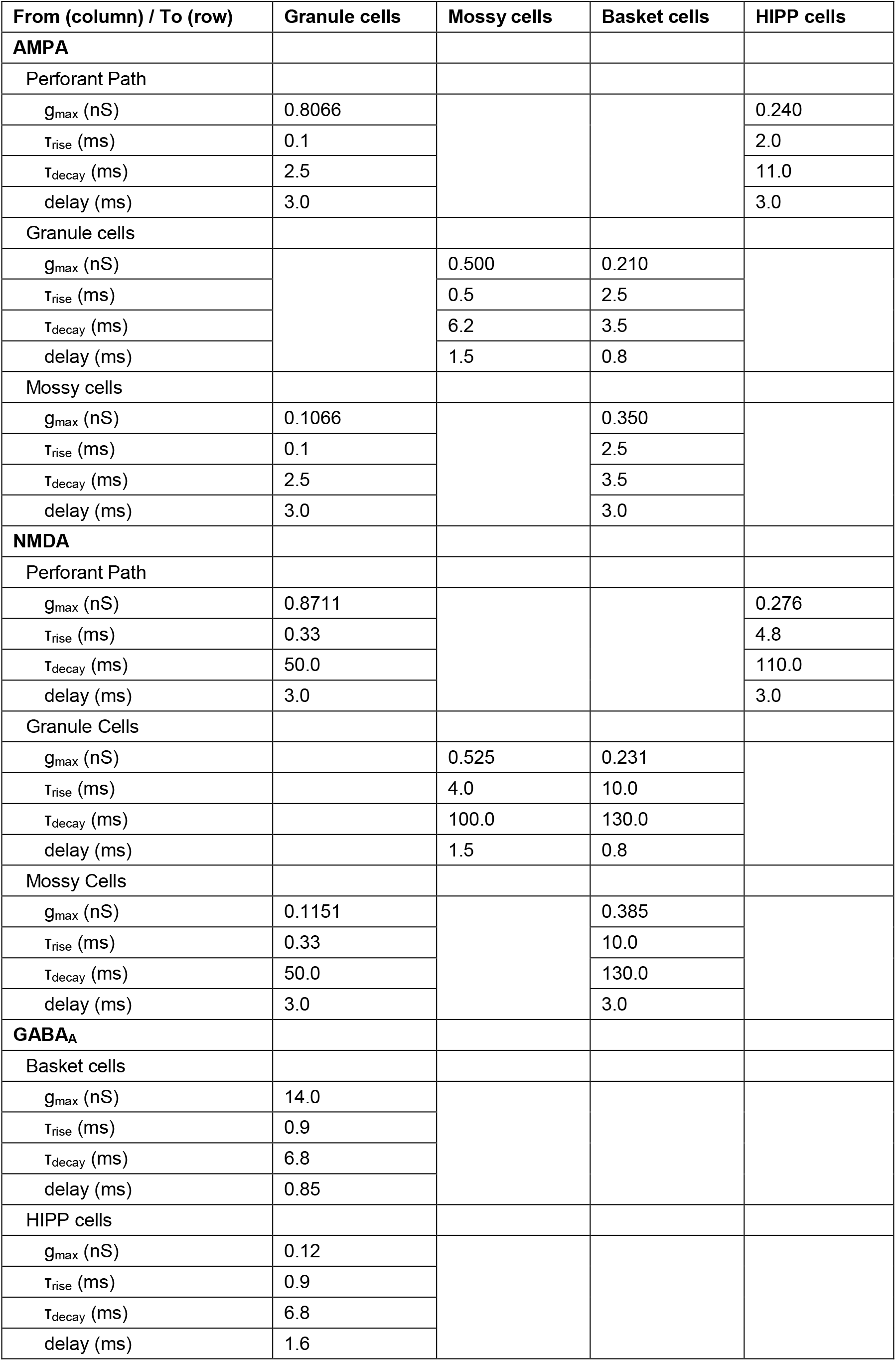
Synaptic parameters of the dentate network

## The DG network model

The DG network model consists of 2000 simulated GCs, a scale that represents 1/500 of the one million GCs found in rat brains (West et al., 1991). The chosen number of GCs provides enough power to explore pattern separation, while maintaining computational efficiency. The population of GCs is organized in non-overlapping clusters, with each cluster containing 20 GCs, respectively. This kind of organization roughly corresponds to the lamellar organization along the septotemporal extent of DG (Sloviter and Lømo, 2012).

Apart from the principal excitatory dentate cells (GCs), the model comprises two kind of inhibitory interneurons, the perisomatic (BCs), which form synapses at the soma of the GCs, and dendritic (HIPP) inhibitory cells, which contact the GCs at their distal dendritic compartments. There is one BC per cluster of GCs, which in turn corresponds to 100 simulated BCs in the model. This is a form of “winner-take-all” competition (Coultrip et al., 1992) in which all, but the most strongly activated GCs in a cluster, are silenced. Given 100 clusters in the model, and with one winner within each cluster, approximately 5% of GCs are active for a given stimulus; this is in agreement with the theoretically and experimentally estimation of 2–5% granular activity in the substrate (Treves et al., 2008; Danielson et al., 2016). Moreover, the model includes simulated hilar MCs and HIPP cells. Estimated numbers for these neuronal types vary from 30,000 to 50,000 MCs in rats (West et al., 1991; Buckmaster and Jongen-Rêlo, 1999), which in turn corresponds to 3–5 MCs per 100 GCs. Accordingly, the model includes 80 MCs per 2,000 GCs. Experimental counts for HIPP cells vary significantly, but the latest estimates suggest about 12,000 HIPP cells in rats (Buckmaster and Jongen-Rêlo, 1999) meaning less than 2 HIPP cells per 100 GCs. To reflect this empirical data, we simulated 40 HIPP cells in the network model (Figure 1A).

External input to the network model is provided by 400 afferents representing the major input that DG receives from Entorhinal Cortex (EC) Layer II cells, via the Perforant Path (PP). The ratio of GCs to PP afferents is aligned with estimations of about 200,000 EC Layer II cells in the rat (Amaral et al., 1990), suggesting a ratio of 20 EC cells per 100 GCs. Therefore, the model incorporates synaptic input that corresponds to 400 EC Layer II cells. For simplicity, the input cells are simulated as independent Poisson spike trains, with frequency of 40 Hz, which is in line with experimental data (Hafting et al., 2005). Previous experimental studies have shown that dentate GCs receive input from 10% of the 4,000 afferents that contact a given GC in the rat during a task (McNaughton et al., 1991), which in turn suggests that an approximate 10% of EC Layer II cells are active. The simulations reported here assume that 10% is the active PP afferents representing a given stimulus. According to McNaughton et al., 1991, 10% of the total entorhinal input is necessary to discharge one GC. However, the EC-GC connection is sparse, with each GC receiving input from about 2% of EC Layer II neurons. Assuming only 400 input cells; one GC could have only 8 afferents from EC, which in turn would make it impossible for the GC to become active. As a compromise, we used a randomly determined 20% of EC Layer II cells as input to each GC and additionally, 20% randomly determined EC Layer II cells as input to each HIPP cell; GCs contact each MC with 20% probability; GCs and HIPP cells each feedback to contact a randomly determined 20% of GCs and finally, each MC connects with every BC in the network. Connections are initialized randomly (uniform random distribution) before the start of the simulations and remain fixed across all simulations (no rewiring). The connectivity matrix was the same for all experiments and across all using GC models, apart from the PP➔GC, and HIPP➔GC synapses due to the difference in GC number of dendrites.

## Pattern Separation and Data Analysis

Generally, a network performs pattern separation whenever the similarity between two distinct input patterns is higher than the similarity between the corresponding output patterns (Figure 1B). In this work, the input patterns are presented as the activity along the 400 PP afferents. Each input pattern has 40 active PP afferents (10% input density), an amount of which are common between two patterns; hence the two patterns have a degree of similarity. Active cell is considered every GC that fires at least one spike during stimulus presentation (Myers and Scharfman, 2009), thus the output patterns correspond to the active GCs. In order to quantify the pattern separation efficiency we used a metric denoted by *f*_1_ (‘population distance’):

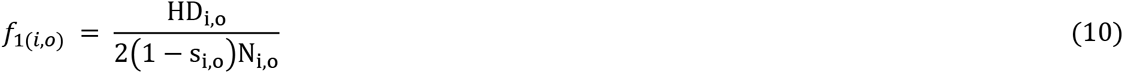

where the *i* and *o* subscripts denote input and output, respectively, *s* denotes the sparsity (i.e., the ratio of silent neurons to all neurons), N the number of neurons, and HD is the hamming distance between two binary patterns (Hamming, 1950), defined as the number of positions at which the corresponding values are different. The factor of 2 in the denominator is used to limit our distance measure at zero. Our network is said to perform pattern separation if the input distance is smaller than the output distance, i.e., when *f*_1(*i*)_ < *f*_1(*o*)_. Thus far, the distance between two binary patterns is calculated using only the HD metric. Although we construct the input patterns to have the same sparsity (i.e., 10% are active), the corresponding output patterns do not necessarily have the same activity level. As we want to examine the differences among the active neurons of each pattern we disengage the dependence on sparsity by dividing the HD with the number of neurons that are active in a pattern. In our case, the output patterns are vectors, each with length equal to 2000 and 2–5% active neurons in 12 dendrites case, which in turn correspond to 40–100 neurons. The total number of active neurons lies in the range of 40–200. For example, if the HD between two patterns is 20, the old metric gives a distance equal to 0.01 whereas the *f*_1_ metric ranges from 0.10–0.25, depending on the percentage of active GC neurons. Thus, using the proposed *f*_1_ metric, differences only between active neurons are taken into account, making the metric more robust across different levels of sparsity.

We constructed four groups of input pattern pairs, with different degrees of similarity and calculated the input and the corresponding output population distances for each group independently. Firstly, we constructed a variety of input patterns with input density 10% (i.e., 40 active neurons) and consequently, four additional input patterns were built with 40 active neurons, 8, 16, 24 and 32 of which are common between patterns, respectively, which, in turn, corresponds to *f*_1(*input*)_ = 0.4,0.3,0.2 and 0.1. The reasoning behind this approach is to examine highly overlapping patterns *f*_1(*input*)_ = 0.1,0.2), as well as less similar ones *f*_1(*input*)_ = 0.3,0.4). Thus, each trial was composed of two simulations using two input patterns within each group.

Whereas the *f*_1_ metric quantifies the distance between two binary vectors containing active and non-active neurons (‘population distance’), we used an additional metric, denoted by *f*_2_, which quantifies the distance in the firing rates of common neurons that encode two patterns by using their firing rates (‘rate distance’) (Figure 1C). The *f*_2_ metric is calculated by dividing, for each neuron that is active in both patterns, its mean firing rate given one stimulus (40 Hz) by its mean firing rate given a stronger stimulus (50 Hz), and averaging these ratios across the population of input and output neurons, respectively (Leutgeb et al., 2004). We subtract this ratio from one in order to convert the ‘rate similarity’ into a ‘rate distance’:

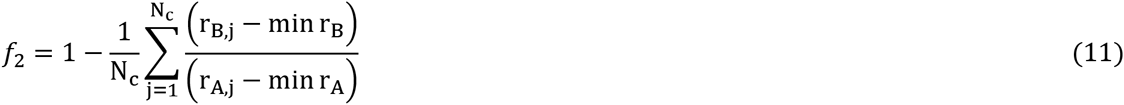

where N_c_ denotes the number of common neurons that are active for bot inputs, r the firing rate of *j-th* neuron using input A and B, respectively. Here, B represents the low firing frequency input, while A the high frequency input. We subtract the global minimum firing rate of GCs found in all trials in order to normalize the dynamic range of firing rates.

For this experiment, the population of active EC neurons in each pair of inputs was identical. The network performs pattern separation if the input ‘rate distance’ is smaller than the corresponding distance in the output, i.e., *f*_2(*input*)_ < *f*_2(*output*)_. In each trial, the two stimuli were presented to the network and their ‘rate distance’, both in input and output, was estimated. The results were then averaged across all trials (50 trials for each model).

For each trial, the network was simulated for *850* milliseconds (ms). The first *300* ms were simulated in order for the network to reach its stable state, so they were excluded from the analysis. The input onset was at *300* ms and the stimulus was applied for *500* ms. The last *50* ms were simulated in order for the network to reach again its steady state and they were excluded from the analysis as well. The time step for all simulations was set to *0.1* ms.

The data analysis and the figures describing the results were made using custom made programs in python2.7.10™ (<u>www.python.org)</u> while the statistical analysis was made using the R3.3.1 programming language (<u>https://www.r-project.org</u>). We used the two-sided, two-sample Wilcoxon signed-rank test for the pattern separation efficiency comparison and the two-sided, two-sample Kolmogorov-Smirnov test (K-S test) to compare the GC activity probability density functions (Neuhäuser, 2011). The model will be available for download at <u>http://dendrites.gr/en/publications-8/software-23</u>.

## Acknowledgements

This research was supported by the European Research Council Starting Grant ‘dEMORY’, (ERC-2012-StG-311435). We would like to thank Alessandro Treves, Idan Segev and Nelson Spruston for their valuable feedback on the project. In addition, we would like to thank Athanasia Papoutsi and Panagiotis Bozelos for their helpful comments on the manuscript, as well as George Kastellakis for his assistance in the software implementation and Pavlos Pavlidis for his guidance on the statistical analysis and finally, all members of the Poirazi lab for their comments during the various phases of the project.

